# Multiple Pol determinants constrain koala retrovirus A replication

**DOI:** 10.64898/2026.10.13.759007

**Authors:** Milky Abajorga, Ann Dauphin, Tianxiong Yu, Jeremy Luban

## Abstract

Koala retrovirus subtype A (KoRV-A) is a gammaretrovirus that occurs in both endogenous and horizontally transmissible forms in wild koala populations. KoRV infection is associated with neoplasia and increased susceptibility to opportunistic infections. Its ongoing incorporation into the koala germline provides a rare contemporary example of retroviral endogenization in a mammalian host. We generated a consensus KoRV-A clone using viral sequences from an infected wild koala and compared it with a published clone representing a viral isolate from a captive koala. Both replicated far less efficiently than Moloney murine leukemia virus (Mo-MLV), the best-characterized gammaretrovirus, during spreading infection in human cell lines, prompting us to investigate the viral features underlying this difference. Following transfection, KoRV-A and Mo-MLV produced comparable levels of viral RNA and intracellular Gag and Pol proteins, yet KoRV-A yielded approximately 1,000-fold less particle-associated reverse transcriptase (RT) activity and markedly lower amounts of Gag-and Pol-derived proteins in released virions. Reduced accumulation of extracellular viral proteins persisted with protease-inactive precursors, indicating that impaired proteolytic maturation alone could not explain this phenotype. KoRV-A/Mo-MLV chimeras showed that particle-associated RT activity depended on both RT sequence and compatibility between protease (PR) and the cleavage sites flanking RT; replacement of the entire KoRV-A PR-RT region with Mo-MLV sequence restored RT activity to near Mo-MLV levels. Transmission electron microscopy (TEM) and spatial analysis revealed that KoRV-A virions accumulated in substantially larger and denser extracellular clusters than Mo-MLV virions. RT and PR–RT chimeras exhibited increased particle abundance and apparent clustering, although most spatial differences were attenuated after accounting for particle density. Whether extracellular particle aggregation contributes directly to inefficient replication remains unclear. Together, these findings identify coordinated interactions among PR, RT, and their cleavage-site context as determinants of KoRV-A enzymatic activity and reveal an unusual extracellular particle organization associated with a gammaretrovirus.

**Author Summary:** Retroviruses insert a DNA copy of their genome into the chromosomes of infected cells. When this occurs in germ cells, the integrated viral DNA can be transmitted to offspring and, over generations, may become a permanent part of the host species’ genome. Koala retrovirus A (KoRV-A) is unusual because this process is occurring in koalas today, allowing us to study retroviral endogenization as it happens rather than reconstructing it from ancient viral remnants. We compared KoRV-A with Moloney murine leukemia virus (Mo-MLV), the best-studied gammaretrovirus. We found that KoRV-A spread much less efficiently in human cell cultures and produced approximately 1,000-fold less reverse transcriptase (RT) activity. KoRV-A also released substantially lower amounts of viral proteins, and virions accumulated in large groups outside cells. By exchanging regions between the two viruses, we determined that RT activity depended on both the enzyme itself and its compatibility with the protease (PR) and PR cleavage sites. These experiments mapped several of these properties to distinct parts of the viral *pol* gene. Thus, multiple barriers constrain KoRV-A replication and provide insight into a retrovirus currently invading a mammalian germline.

## Introduction

Retroviruses are enveloped RNA viruses that spread by reverse transcribing their RNA genomes into DNA and integrating this cDNA into the host genome [1,2]. Their genomes encode three major genes: gag, which produces the structural proteins of the virion [3–8]; pol, which encodes the viral enzymes protease (PR), reverse transcriptase (RT), and integrase (IN) [9–15]; and env, which encodes the envelope glycoproteins responsible for receptor recognition and membrane fusion [16–21]. The Gag polyprotein is cleaved into matrix (MA), capsid (CA), and nucleocapsid (NC), and, depending on the retroviral lineage, additional proteins or spacer peptides. MA associates with the inner leaflet of the viral membrane [3–5], CA forms a protein lattice that encloses the diploid positive-sense RNA genome [3,6], and NC binds and packages the virion genomic RNA [7,8]. Through translational readthrough of the gag termination codon, gammaretroviruses synthesize the Gag–Pol fusion polyprotein, which is subsequently processed by the viral protease into its mature enzymatic components [22,23]. Efficient viral replication therefore depends on coordinated interactions among structural proteins, Pol maturation, and Env-mediated entry.

Murine leukemia virus (MLV) has served as a principal model for understanding gammaretrovirus replication, virion assembly, and maturation [24]. Although endogenous MLV elements are abundant in mice, they entered the germline millions of years ago and have accumulated mutations that render most copies replication defective [25–29]. Hence, MLV is a powerful experimental model for retroviral biology, but offers limited insight into the earliest stages of retroviral germline invasion.

Koala retrovirus (KoRV) is one of the few known retroviruses currently undergoing endogenization in a mammalian host. KoRV-A, the transmissible subtype responsible for germline invasion, is present in nearly all northern Australian koalas but remains incompletely endogenized in southern populations [30,31]. This ongoing invasion provides a rare opportunity to investigate the molecular features of a retrovirus during active germline colonization. In addition to vertical transmission, KoRV can spread horizontally, and infection has been associated with neoplasia, immunological dysfunction, and increased susceptibility to infectious disease [32–37]. While several of these disease associations are linked predominantly to exogenous KoRV subtypes such as KoRV-B, KoRV-A is the subtype responsible for germline transmission [30,36–39].

Considerable effort has focused on understanding host responses to KoRV infection, including piRNA-mediated silencing in the germline, innate immune signaling pathways, and adaptive immune responses directed against Env [40–44]. However, the intrinsic replication biology of KoRV-A itself remains poorly understood. For example, why KoRV-A replicates inefficiently relative to other gammaretroviruses or whether viral enzymatic activity, virion maturation, particle production, entry, or extracellular organization contribute to this phenotype is unknown. To address these questions, the replication properties of KoRV-A were compared with those of

Moloney MLV (Mo-MLV). KoRV-A expressed viral RNA and intracellular viral proteins at levels comparable to Mo-MLV, but demonstrated reduced particle-associated RT activity, inefficient spread, and extensive extracellular virion clustering. Using a series of KoRV-A/Mo-MLV chimeric viruses, quantitative spatial analysis of virion organization, characterization of viral protein expression and processing, and electron microscopy, we identified coordinated Pol-domain and cleavage-site requirements that constrain RT activity.

## Materials and Methods

### Sequence alignment and sliding-window identity analysis

KoRV-A and Mo-MLV protein sequences were aligned using the MUSCLE Alignment tool with the PPP algorithm in Geneious Prime version 2026.1.2 [45]. The resulting amino-acid alignments were exported with gap characters retained. Sequence alignments were visualized using ESPript to display residue conservation and insertion/deletion positions [46].

Sliding-window amino-acid identity was calculated using a custom Python script. A 50-amino-acid window was advanced across each alignment in one-residue increments. Within each window, percent identity was calculated as the number of alignment columns containing identical, non-gap amino acids divided by the number of evaluated columns, multiplied by 100. Columns containing a gap in only one sequence were treated as nonidentical, whereas columns containing gaps in both sequences were excluded from the denominator. Identity values were plotted against the center position of each window using one-indexed alignment coordinates.

### Generation of the KoRV-A consensus sequence

A consensus sequence representing transcriptionally active KoRV-A was generated from liver RNA-sequencing reads obtained from koala K63855 [41]. RNA-sequencing reads aligning to ribosomal RNA were removed using Bowtie2 v2.2.5 with default parameters [47]. The remaining reads were aligned to transposon consensus sequences using HISAT2 v2.1.0 with default parameters [48]. Reads corresponding to the KoRV-A reference sequence (GenBank accession AF151794) were assembled using SPAdes with default parameters [49,50]. The resulting KoRV-A sequence was synthesized and cloned into the pBluescript II SK(+) vector.

### Plasmids

All plasmids used in this paper are described in S1 Table. Plasmids were synthesized by GENEWIZ using their PriorityGENE service or generated using FastDigest restriction-enzyme cloning (Thermofisher Scientific), Gibson Assembly (New England Biolabs; cat. no. E2611), In-fusion cloning (Takara Bio; cat. no. 638946), or PCR-based site-directed mutagenesis. Gene blocks and primers utilized for cloning were obtained from GENEWIZ or Integrated DNA Technologies (IDT). Complete plasmid sequences and maps have been deposited in the Dryad Digital Repository under the reserved DOI: https://doi.org/10.5061/dryad.cjsxksnpj. Physical plasmids will be made available through BEI Resources (accession numbers pending).

### Tissue culture

HEK293T cells (ATCC CRL-3216) and Jurkat cells (ATCC TIB-152) were obtained from the American Type Culture Collection (ATCC) and maintained in Dulbecco’s Modified Eagle Medium (DMEM) (Gibco; cat. no. 11955-065) and RPMI 1640 Medium (Gibco; cat no. 11875-093), respectively, supplemented with 10% heat-inactivated cosmic calf serum (CCS) (Cytiva; cat. no. SH30087.03), 1% sodium pyruvate (Corning; cat. no. 25-000-CI), 1× GlutaMAX (Gibco; 35050-061), 1% MEM nonessential amino acids (Corning; cat. no. 25-051-CI), and 1% HEPES buffer (Corning; cat. no. 25-060-CI). Cells were cultured at 37°C with 5% carbon dioxide (CO_2_) and monitored for mycoplasma contamination using the Mycoplasma Detection kit (Lonza LT07-318).

### Transfection

All transfections were done using TransIT-LT1 (Mirus Bio; cat. no. MIR 2306) according to the manufacturer’s instructions. Cells were plated at 60% confluency and transfected with plasmids. For normalization of transfection efficiency, cells were cotransfected with the EGFP retrovirus expression plasmids and pEF1*α*-E2 Crimson (Takara; cat. no. 631981), an E2 Crimson expression plasmid, at a 5:1 mass ratio.

### Transduction

30,000 HEK293T cells or 45,000 Jurkat cells were infected by spinoculation with 10^5^ RT units of either KoRV-A, Mo-MLV, and other retroviral chimeras at 1300×g for 2 h on a table top Eppendorf Centrifuge 5425. The supernatant was removed and the cells were resuspended in 700 µL of the appropriate culture medium. Cells were plated in a 24-well plate and viral supernatant was collected every 2-4 days for RT assay. Cells were split in half every other day once confluent. Cells and viral supernatant were collected for western blot as well.

### Exogenous Reverse Transcriptase assay

A one-step PCR based assay was used to quantify virion-associated RT activity of virions [51]. Supernatant (5 µL) was lysed in an equal volume of 0.25% Triton X-100, 50 mM potassium chloride (KCl), 100 mM Tris(hydroxymethyl)aminomethane hydrochloride (Tris–HCl; pH 7.4), 40% glycerol, and RiboLock RNAse inhibitor (0.4 U/μL; ThermoFisher; cat. no. EO0382). Viral lysate was diluted 1:100 in a buffer containing 5 mM ammonium sulfate [(NH_4_)_2_SO4], 20 mM KCl, and 20 mM Tris–HCl pH 8.3. Diluted viral lysate (10 µL) was combined with 10 µL of reaction mix containing 10 mM (NH_4_)_2_SO4, 40 mM KCl, 40 mM Tris–HCl pH 8.3, 10 mM magnesium chloride (MgCl_2_), 0.2 mg/mL bovine serum albumin (BSA), 1/10,000 SYBR Green I (Invitrogen), 400 μM dNTPs, 1 μM forward primer (TCCTGCTCAACTTCCTGTCGAG), 1 μM reverse primer (CACAGGTCAAACCTCCTAGGAATG), 8 μg/mL MS2 RNA (Roche), RiboLock RNAse inhibitor (0.04 U/μL), and Hot Start Polymerase (5 U/mL; Promega; cat. no. M5001). The reaction was run on a Bio-Rad CFX96 Touch Real–Time PCR Detection System using the following program: 42°C for 20 min, 95°C for 2 min, and 40 cycles of 95°C for 5 s, 72°C for 15 s, and fluorescence acquisition at 80°C for 5 s. RT activity was quantified using a standard curve generated from the viral supernatant of cells transfected with the HIV *gagpol* expression plasmid, psPAX2. A no-virus control, in which no retrovirus-containing sample was added, was included to confirm that amplification depended on virion-associated RT activity rather than contaminating DNA or nonspecific amplification.

### Production of rabbit antisera targeting KoRV-A proteins

Polyclonal antibodies against capsid (CA), matrix (MA), reverse transcriptase (RT), and integrase (IN) of KoRV-A were generated by the Pocono Rabbit Farm and Laboratory Inc using the 49 Day Protocol. Peptides were selected using the AbDesigner program (NHLBI) to optimize immunogenicity and to be distinguishable from MLV peptides [52]. Rabbits were immunized with the peptides TEERERRRDRRQEKNC, RSRRPRSPTGDTGPDC, KKPGTNDYRPVQDLR, and TYRESGRRQRGDRPG for KoRV-A CA, MA, RT, and IN, respectively.

### Affinity purification of antibodies

One mL of packed Affi-Gel 10 (Bio-rad; cat. no. 1536099) in a polypropylene column (Biorad; cat. no. 7311550) was washed with 3 bed volumes of ddH_2_O and equilibrated with 3 bed volumes of 0.1 M HEPES pH 7.4. Five mg of peptide resuspended in 1 mL of 0.1 M HEPES pH 7.4 was incubated with the affi-gel column for 4 h at 4°C. Conjugation was terminated with 100 µL of 1 M ethanolamine pH 8.0 by rocking gently for 1 h at 4°C. After collection of the flowthrough, the affi-gel column was washed with ddH_2_O, then 1× PBS (Phosphate Buffered Saline), followed by 0.1 M glycine pH 2.3, and finally with 1× PBS again. Conjugated beads were incubated with 5 mL of serum in 21 mL of 1× PBS in a falcon tube overnight with rocking. Resin was transferred to the column, flowthrough was collected, and resin was washed with 1× PBS 3 times. Eight 1 mL fractions of antibody were collected by eluting with 0.1 M glycine pH 2.3 into a tube with 100 μL of 1 M Tris pH 9.5. Antibody fractions were diluted by 50% by adding glycerol and sodium azide was added to a final concentration of 0.02%. Antibody fractions were concentrated using an Amicon Ultra Centrifugal Filter, with a 50 kDa cut-off (Millipore Sigma; cat. no. UFC505008).

### Western blot

Cells were trypsinized and washed with 1× PBS. Equal numbers of cells were lysed in RIPA buffer composed of 50 mM Tris-HCl (pH 7.4), 150 mM NaCl, 1% NP-40, 0.5% sodium deoxycholate, and 0.1% SDS. The RIPA buffer was supplemented with 1× Halt™ Protease Inhibitor Cocktail, EDTA-Free (Thermo Scientific; cat. no. 78439) and Pierce™ Universal Nuclease for Cell Lysis (Thermo Scientific; cat. no. 88702). Cell lysates were mixed with either 2× Laemmli Sample Buffer (Bio-Rad; cat. no. 1610737) or 4× Laemmli Sample Buffer (Bio-Rad; cat. no. 1610747), supplemented with β-mercaptoethanol. Samples were loaded as equal cell equivalents.

Viral particles in culture supernatants were pelleted by ultracentrifugation at 120,000 × *g* for 1 h at 4°C using a TLA-55 rotor in a Beckman Coulter Optima MAX-XP ultracentrifuge. Viral pellets were suspended by adding 2× Laemmli Sample Buffer supplemented with β-mercaptoethanol directly to each pellet. Cell-lysate and viral-pellet samples were heated at 95°C for 5 min before electrophoresis.

Proteins were separated by SDS-PAGE using 4–20% Mini-PROTEAN TGX Precast Gels (Bio-Rad; cat. nos. 4561096, 4561094, and 4561095). Proteins were transferred to nitrocellulose membranes (Bio-Rad; cat. no. 1620112) by wet transfer at 100 V for 1 h and 45 min. An ice pack was placed in the transfer tank, and the apparatus was surrounded by ice to minimize heating during transfer. Membranes were blocked in Intercept TBS Blocking Buffer (LI-COR Biosciences; cat. no. 927-60001) for 1 h at room temperature with agitation. Membranes were subsequently incubated overnight (approximately 16 h) at 4°C on a nutator with primary antibodies diluted in Intercept TBS Blocking Buffer.

Primary antibodies included anti-FLAG epitope (1:1,000; Novus Biologicals; cat. no. NB600-345), anti-GFP (1:1,000; Takara Bio; cat. no. 632381), anti-β-actin (1:1,000; Abcam; cat. no. ab8226), anti-KoRV-A CA, anti-KoRV-A MA, anti-KoRV-A RT, anti-KoRV-A IN, and anti-MLV CA (clone R187). Membranes were washed three times for 5 min each with 1× TBS-T containing 0.1% Tween-20 and incubated for 1 h at room temperature on a rocker with secondary antibodies diluted in Intercept TBS Blocking Buffer. Secondary antibodies included goat anti-rabbit IgG (1:10,000; LI-COR Biosciences; cat. no. 926-32211), goat anti-rat IgG (1:10,000; LI-COR Biosciences; cat. no. 925-32219), and goat anti-mouse IgG (1:10,000; LI-COR Biosciences; cat. no. 926-68070). Membranes were then washed three times for 5 min each with 1× TBS-T and imaged using an Odyssey CLx Imaging System (LI-COR Biosciences).

### RT-qPCR

Total RNA was isolated from cells using the RNAzol Reagent (Molecular Research Center; cat. no. RN 190) in accordance with the manufacturer’s instructions. Residual genomic DNA was removed using the TURBO DNA-free Kit (Invitrogen; cat. no. AM1907). First-strand cDNA was synthesized using the iScript Advanced cDNA synthesis kit (Bio-Rad; cat. no. 1725038) in a Bio-Rad thermal cycler. qPCR was performed using iTaq Universal Sybr Green Supermix (Bio-Rad; cat. no. 1725121) on a Bio-Rad CFX96 Real-Time PCR Detection System for 40 cycles of 95°C for 5 s and 60°C for 30 s. Primer sequences are provided in S2 Table.

### Flow cytometry and analysis

Cells were trypsinized and washed twice with FACS buffer (1× PBS, 0.1% sodium azide, and 1% heat-inactivated CCS), fixed in BD Cytofix Fixation Buffer (BD Biosciences; cat. no. 554655) for 30 min at 4°C, washed twice with FACS buffer, and resuspended in FACS Buffer.

For analysis of cells expressing EGFP alone, data were acquired with the BD Accuri C6 flow cytometer, with at least 100,000 events collected within the primary cell population. Data were exported as FCS files and analyzed using FlowJo (version 10.10.0). Cellular debris was excluded based on forward scatter (FSC) and side scatter (SSC) properties. Singlet cells were identified by FSC-A versus FSC-H gating. EGFP-positive cells were identified using FITC-H fluorescence, with the fluorescence threshold established using an EGFP-negative population. The same gating strategy and fluorescence threshold were applied to all samples within each experiment.

For samples expressing both E2-Crimson and EGFP, data were acquired on a BD FACSymphony A5 cytometer using FACSDiva software (version 9.3.1). Compensation was applied uniformly to all samples within an experiment using mock-transfected cells as the unstained control and cells expressing E2-Crimson alone as single-color compensation control. Compensated data were exported as FCS files and analyzed using FlowJo (version 10.10.0). The same sequential gating strategy was applied to all samples within each experiment. Gates defining E2-Crimson-positive and EGFP-positive populations were established using the corresponding negative controls and then applied consistently across samples. To normalize GFP expression to the cotransfected E2-Crimson reporter, analysis was restricted to cells positive for both GFP and E2-Crimson. Within this double-positive population, GFP median fluorescence intensity (MFI) was divided by E2-Crimson MFI and multiplied by 100. The resulting normalized GFP MFI was used for subsequent analyses.

### Transmission Electron Microscopy (TEM)

At 36 hours post-transfection, HEK293T cells were fixed by removing half of the culture medium and replacing it with an equal volume of fixative containing 2.5% glutaraldehyde and 1.6% paraformaldehyde in 0.1 M sodium cacodylate buffer (pH 7.2). After a 10 min incubation at room temperature, the fixative-media mixture was removed and replaced with fresh fixative, and cells were incubated for an additional 1 h at room temperature. Samples were washed three times for 10 min each in 0.1 M sodium cacodylate buffer (pH 7.2) and post-fixed in 1% osmium tetroxide in ddH_2_O for 1 h at room temperature. Following three 10-min washes in ddH_2_O, cells were gently scraped from the culture plate with a soft plastic spatula and transferred to microcentrifuge tubes. Cell pellets were washed three additional times for 10 min each in ddH_2_O and dehydrated through a graded ethanol series (10%, 30%, 50%, 70%, 85%, and 95% for 20 min each), followed by three changes of 100% ethanol. Samples were infiltrated with two changes of 100% propylene oxide, followed by overnight incubation in a 1:1 mixture of propylene oxide and SPI-Pon 812 resin. Approximately 24 h later, cell pellets were transferred through four changes of freshly prepared SPI-Pon 812R hard-formulation epoxy resin comprising DDSA (catalog no. 02827-AF), Araldite 502 epoxy resin (catalog no. 02816-AB), SPI-Pon 812R epoxy resin (catalog no. 02659R-AB), and DMP-30 epoxy accelerator (catalog no. 02823-DA; all from SPI Supplies Division, Structure Probe, Inc.). Samples were embedded in tubes containing the same resin formulation and polymerized at 70°C for 48 h. Epoxy blocks were trimmed, and 70-nm ultrathin sections were cut on a Reichart–Jung ultramicrotome using a diamond knife. Sections were collected onto 200-mesh copper support grids (SPI; cat. no. 2620C) and contrasted with 2% lead citrate and 4% uranyl acetate. Samples were examined using a Philips CM10 transmission electron microscope operated at an accelerating voltage of 100 kV. Grids were surveyed to locate extracellular virions, and images were acquired when a field containing at least one virion was identified using a Gatan Erlangshen ES1000W TEM CCD camera and DigitalMicrograph software (version 3.11.2) for GMS 1.6.2. Accordingly, the analyzed images represent selected virion-containing fields rather than randomly sampled fields across the entire grid.

### Spatial analysis of virion clustering by nearest-neighbor and DBSCAN analysis

An automated blob-detection workflow was initially evaluated for identification of extracellular virions in TEM images. However, the automated approach produced both false-positive detections and incomplete identification of virions. To minimize these errors, virions in TEM images were manually identified and annotated in ImageJ2 (2.16.0/1.54p; National Institutes of Health) based on particle size, morphology, and electron density. The x-and y-coordinates of each virion centroid were recorded and exported as CSV files. Quantitative spatial analyses were restricted to TEM images acquired at a single magnification (92,000×) and covering the same physical field of view, allowing direct comparison of virion counts and spatial measurement across images. Spatial analyses were performed using custom scripts written in Python (version 3.9.6), with pandas (version 2.3.3), NumPy (version 2.0.2), SciPy (version 1.13.1), and scikit-learn (version 1.6.1). For each image, virion centroid coordinates were imported from the corresponding CSV file for downstream analysis.

For images containing two or more virions, pairwise Euclidean distances between virion centroids were calculated. For each virion, the distance to its nearest neighboring virion was determined. The median nearest-neighbor distance and the percentage of virions within 100 nm of another virion were then calculated for each image. Median nearest-neighbor distance was undefined for images containing a single virion and was therefore recorded as not applicable (NaN). Images containing no virions were excluded from spatial analysis.

Virion clustering was assessed using density-based spatial clustering of applications with noise (DBSCAN), with a neighborhood radius (ε) of 100 nm and a minimum of three virions required to define a cluster (min_samples = 3). For each image, the percentage of virions assigned to clusters, the number of clusters, the mean number of virions per cluster, and the number of virions in the largest cluster were calculated for each image. Images containing one or two virions were retained and were assigned values of 0% clustered virions, zero clusters, zero mean cluster size, and zero largest cluster size because they could not satisfy the predefined DBSCAN criterion for a cluster.

Measurements were obtained for each image and then averaged within each biological replicate. A biological replicate was defined as an independent transfection performed on a different day. Biological replicates served as the experimental units for statistical analyses. For each biological replicate, the numbers of TEM images analyzed and virions detected, together with the number and percentage of images containing only one or two virions, were recorded (S3 Table). Image-level spatial measurements were averaged within each biological replicate for comparisons among experimental groups. Median nearest-neighbor distance was calculated only for images containing at least two virions; all other spatial metrics included images containing one or two virions as described above

Particle-group-size and nearest-neighbor-distance distributions were also examined descriptively. To identify particle groups of all sizes, including individual virions and pairs, a separate DBSCAN analysis was performed using ε = 100 nm and *min_samples* = 1. Particle groups were categorized as containing 1, 2, 3–5, 6–8, 9–11, or >11 virions (S16 Fig). Nearest-neighbor distances for individual virions were categorized as <50, 50–100, 100–150, 150–200, 200–300, or >300 nm (S17 Fig). Virions in images containing only one particle were excluded from the nearest-neighbor-distance distribution because no nearest-neighbor distance could be calculated. Particle groups and individual nearest-neighbor distances were pooled across images and biological replicates for visualization.

Because measures of spatial clustering can be influenced by the number of virions in an image, image-level regression analyses for virion density were performed as a sensitivity analysis. Each clustering metric was modeled as a function of the experimental group and the natural logarithm of the number of virions per image, with cluster-robust standard errors estimated at the biological replicate level. The coefficient for the experimental group therefore estimates the difference between groups after accounting for differences in virion abundance per image. Density adjusted analyses are presented as a conservative sensitivity analysis because virion abundance could act either as a confounder of the relationship between viral genotype and clustering or as an intermediate consequence of the clustering phenotype itself; the present experiments do not distinguish between these possibilities.

## Statistical Analysis

Area under the curve (AUC) was calculated separately for each biological replicate over the experimental interval shared across all groups using trapezoidal integration. Data analysis, statistical testing, and figure preparation were performed using GraphPad Prism (version 11.0.2) and custom scripts executed in Python (version 3.9.6). Python analyses used NumPy (version 2.0.2), pandas (version 2.3.3), SciPy (version 1.13.1), scikit-learn (version 1.6.1), statsmodels (version 0.14.6), and Matplotlib (version 3.9.4). The statistical test and, where applicable, the multiple-comparisons procedure used for each experiment are specified in the corresponding Results section and figure legend. Unless otherwise indicated, biological replicates were treated as the independent experimental units, and all tests were two-sided. A *p* value < 0.05 was considered statistically significant. Exact *p* values are reported in the figures, with values below 0.0001 reported as *p* < 0.0001.

### Use of artificial intelligence tools

OpenAI Codex (GPT-5-family model) and Anthropic Claude (Haiku 4.5) were used to assist with language editing, manuscript organization, refinement of the text and the development, drafting, and debugging Python code used for data analysis and figure preparation. The authors critically evaluated and revised all AI-assisted content, verified numerical and statistical statements against the underlying data, and reviewed and validated the analysis code and outputs. All final decisions concerning the analyses, interpretations, limitations, and conclusions were made by the authors, who take full responsibility for the accuracy and integrity of the manuscript. Generative AI was not used to generate, modify, or fabricate primary experimental data or research images.

## Results

### Generation of a KoRV-A consensus provirus and sequence comparison with Mo-MLV

A KoRV-A consensus sequence was derived from RNA-sequencing data obtained from koala K63855, an animal harboring endogenous KoRV-A germline insertions [41]. The consensus sequence was synthesized and cloned into the pBluescript II SK(+) vector. This construct, designated as pBS-KoRV-A, shares 99.6% nucleotide identity with the previously reported pKoRV522 with sequence variations distributed throughout the genome [53]. Ecotropic and amphotropic clones of Mo-MLV were also cloned into the pBluescript II SK(+) vector and designated pBS-MLV and pBS-MLV Ampho, respectively. Pairwise sequence comparison between KoRV-A and Mo-MLV revealed moderate conservation. Gag and Pol proteins share 48.5% and 61.9% amino acid identity, respectively (Fig 1A and 1B). Sequence divergence is concentrated in Gag p12 and the IN N-terminal extension domain (Fig 1C and 1D). KoRV-A harbors a 19-amino-acid deletion within the N-terminal extension region relative to Mo-MLV, corresponding to only 15.9% amino acid identity (Fig 1D) — the largest divergence observed across Pol (Fig 1B and 1D). This deletion is also present in gibbon ape leukemia virus (GaLV) and Hervey pteropid gammaretrovirus (HPG), two retroviruses closely related to KoRV-A, suggesting that it is a conserved feature of this retroviral lineage rather than unique to KoRV-A. The p12 region likewise showed marked divergence, with only 25% amino acid identity. Nevertheless, the PPPY late domain is conserved between KoRV-A and Mo-MLV (Fig 1C).

**Fig 1.**
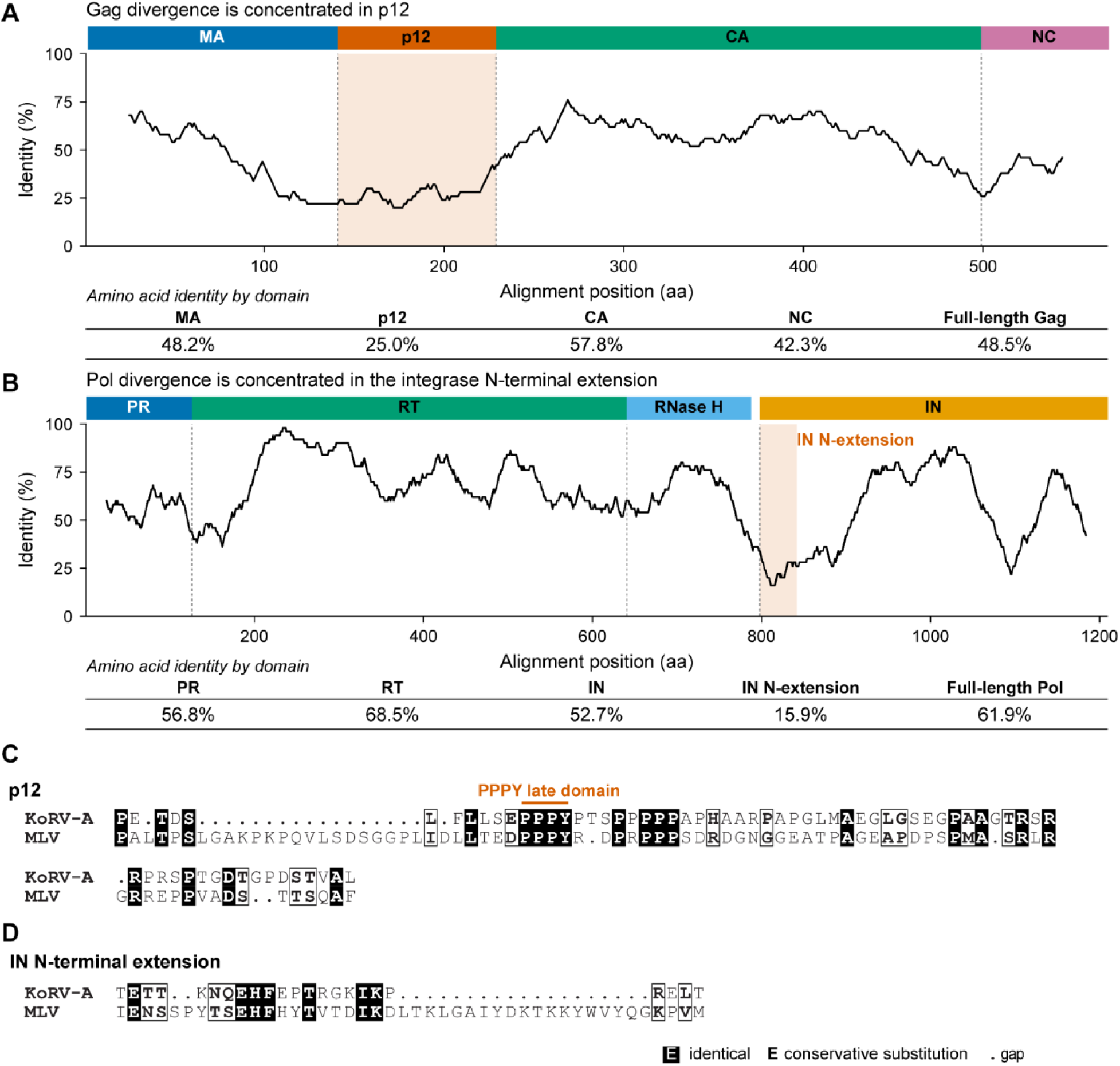
Comparative sequence analysis of KoRV-A and Mo-MLV Gag and Pol proteins. (A) Sliding-window amino acid identity plot across the aligned KoRV-A and Mo-MLV Gag sequences. MA, p12, CA, and NC are indicated above the plot. The table below reports amino acid identity for each Gag domain and full-length Gag protein. (B) Sliding-window amino acid identity plot across the aligned KoRV-A and Mo-MLV Pol sequences. PR, RT, and IN are indicated above the plot. The table below reports amino acid identity for each Pol domain, the IN N-terminal extension, and full-length Pol. (C, D) Pairwise amino acid alignments of the p12 regions (C) and IN N-terminal extension regions (D) of KoRV-A and Mo-MLV. Identical residues are shown as white letters on a black background, conservative substitutions as bold black letters, and alignment gaps as dots.

### KoRV-A replication is inefficient in transformed human cell lines

As an initial characterization of the new constructs, exogenous RT activity was quantified in supernatants from HEK293T cells transfected with pBS-MLV, pBS-MLV Ampho, pKoRV522, pBS-KoRV-A, or the well-established Mo-MLV clone pNCA Ampho. RT activities produced by pBS-MLV and pBS-MLV Ampho transfected cells were comparable to that of pNCA Ampho, suggesting that cloning Mo-MLV into pBluescript II SK(+) did not measurably alter RT activity (Fig 2A). Likewise, cells transfected with pKoRV522 and pBS-KoRV-A produced comparable RT activities, indicating that the synthesized clone recapitulated the low-RT-activity phenotype of the previously published pKoRV522 clone, which was derived from KoRV-A isolated from a captive koala [53]. Both pKoRV522 and pBS-KoRV-A displayed approximately 1,000-fold lower RT activity than the Mo-MLV constructs (Fig 2A).

**Fig 2.**
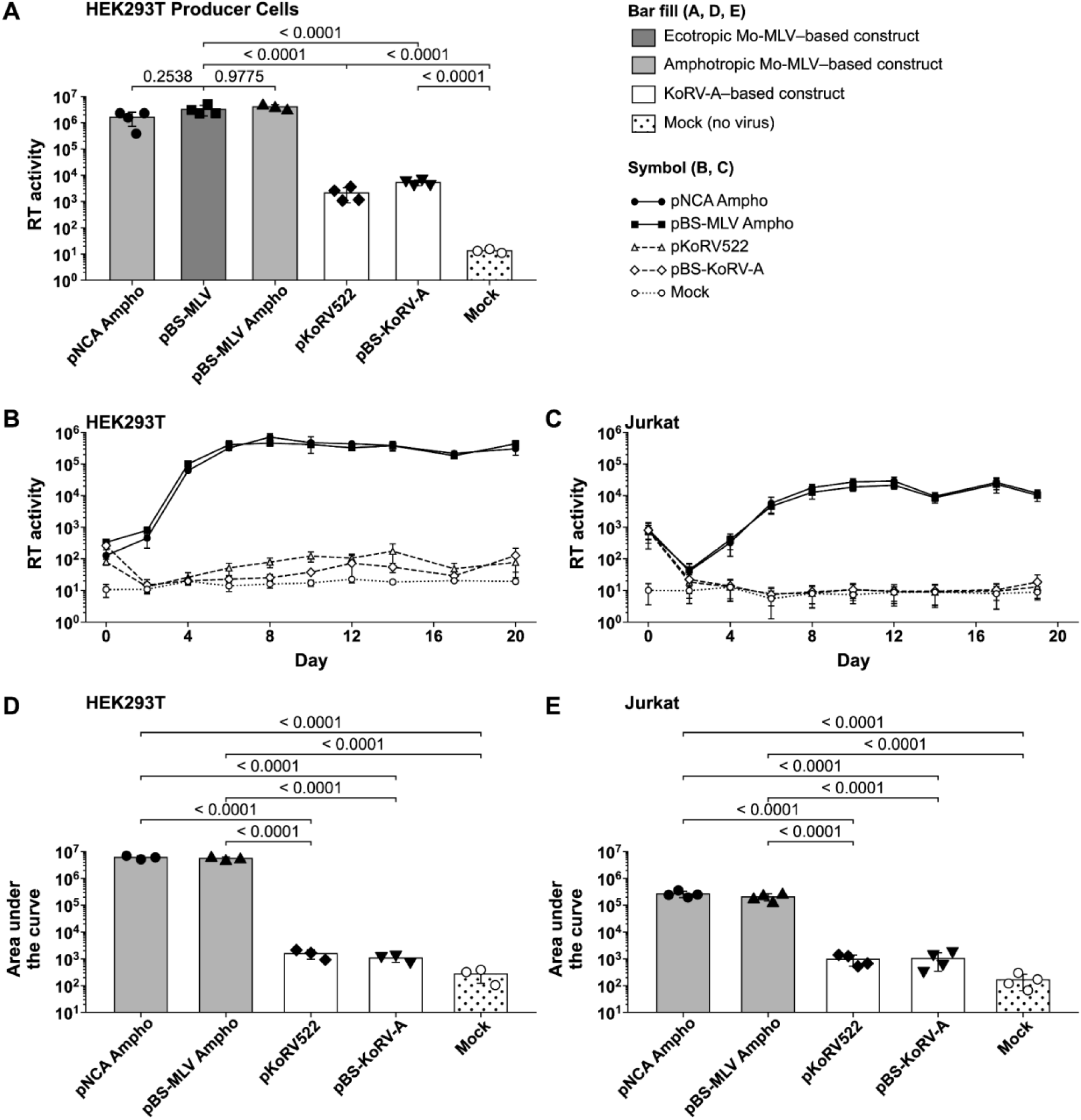
KoRV-A produces lower RT activity and spreads inefficiently in HEK293T and Jurkat cells compared with Mo-MLV. (A) Exogenous RT activity in culture supernatants from HEK293T cells transfected with the indicated KoRV-A or Mo-MLV proviral plasmids. Each point represents an independent biological replicate (n=3-4); data are presented as geometric means ± geometric SD. Tukey-adjusted *p* values from one-way ANOVA are shown. (B, C) Replication kinetics of the indicated viruses in (B) HEK293T cells and (C) Jurkat cells. Cells were spinoculated with virus stock normalized by exogenous RT activity, and viral spread was monitored by measuring exogenous RT activity in culture supernatants at the indicated time points. Values are geometric means ± geometric SD from three independent biological replicates for HEK293T cells and four independent biological replicates for Jurkat cells. (D, E) AUC analysis of replication kinetics in (D) HEK293T cells and (E) Jurkat cells, from (B) and (C), respectively. AUC values were calculated independently for each biological replicate using the trapezoidal rule. Bars show group means ± SD; points represent individual biological replicates. Tukey-adjusted *p* values from one-way ANOVA are shown.

HEK293T and Jurkat cells were then challenged by spinoculation with RT normalized cell-free virus supernatants, and RT activity in culture supernatants was monitored over time. In both cell types, viruses generated from pNCA Ampho and pBS-MLV Ampho exhibited rapidly increasing RT activity before reaching a plateau (Fig 2B and 2C). In contrast, RT activity in cultures inoculated with pKoRV522 and pBS-KoRV-A remained low throughout the experiment. However, a modest increase above the mock background was observed in KoRV-A-inoculated HEK293T cultures (Fig 2B), whereas no appreciable increase was observed in Jurkat cultures (Fig 2C).

Consistent with these replication kinetics, AUC analysis showed markedly lower cumulative RT activity for both KoRV-A clones than for the Mo-MLV controls in HEK293T and Jurkat cells (Fig 2D and 2E). Western blot analysis of CA in matched cell lysates and culture supernatants showed a progressive increase in CA abundance for KoRV-A and Mo-MLV constructs in HEK293T cells (S1 Fig). This finding corroborated the modest increase in RT activity observed in KoRV-A-inoculated cultures and was consistent with low-level KoRV-A replication and spread in HEK293T cells, albeit substantially less efficiently than Mo-MLV. In Jurkat cells, KoRV-A CA was detectable at early time points but declined thereafter, consistent with the absence of a measurable increase in RT activity and indicating that KoRV-A did not undergo detectable spread (S2 Fig). Importantly, normalization of the inocula equated particle-associated RT activity rather than physical particle number and therefore assumes that KoRV-A and Mo-MLV have comparable RT activity per particle. If RT activity per particle differs between the two viruses, the RT-normalized inocula may have contained different numbers of physical particles. This limitation should therefore be considered when interpreting the replication kinetics.

### KoRV-A Gag and Gag–Pol are expressed and proteolytically processed

RT-qPCR using primer pairs targeting *gag*, *pol*, and *env* did not reveal substantial differences in viral transcript abundance between KoRV-A-and Mo-MLV-transfected cells (S3 Fig). Antibodies against MLV CA and KoRV-A CA, MA, RT, and IN were then used to examine variation in viral protein expression in transfected HEK293T cell lysates and virions pelleted from culture supernatants. Because no commercial antibodies against KoRV-A proteins were available, custom polyclonal antibodies were generated in rabbits against peptides corresponding to KoRV-A CA, MA, RT, and IN (see Methods). Western blot analysis detected bands consistent with the full-length Gag precursor, mature CA, mature MA, and additional processing intermediates in transfected cell lysates and pelleted viral supernatants (Fig 3; S4 and S5 Figs). These findings demonstrate the expression and processing of Gag for both retroviruses.

**Fig 3.**
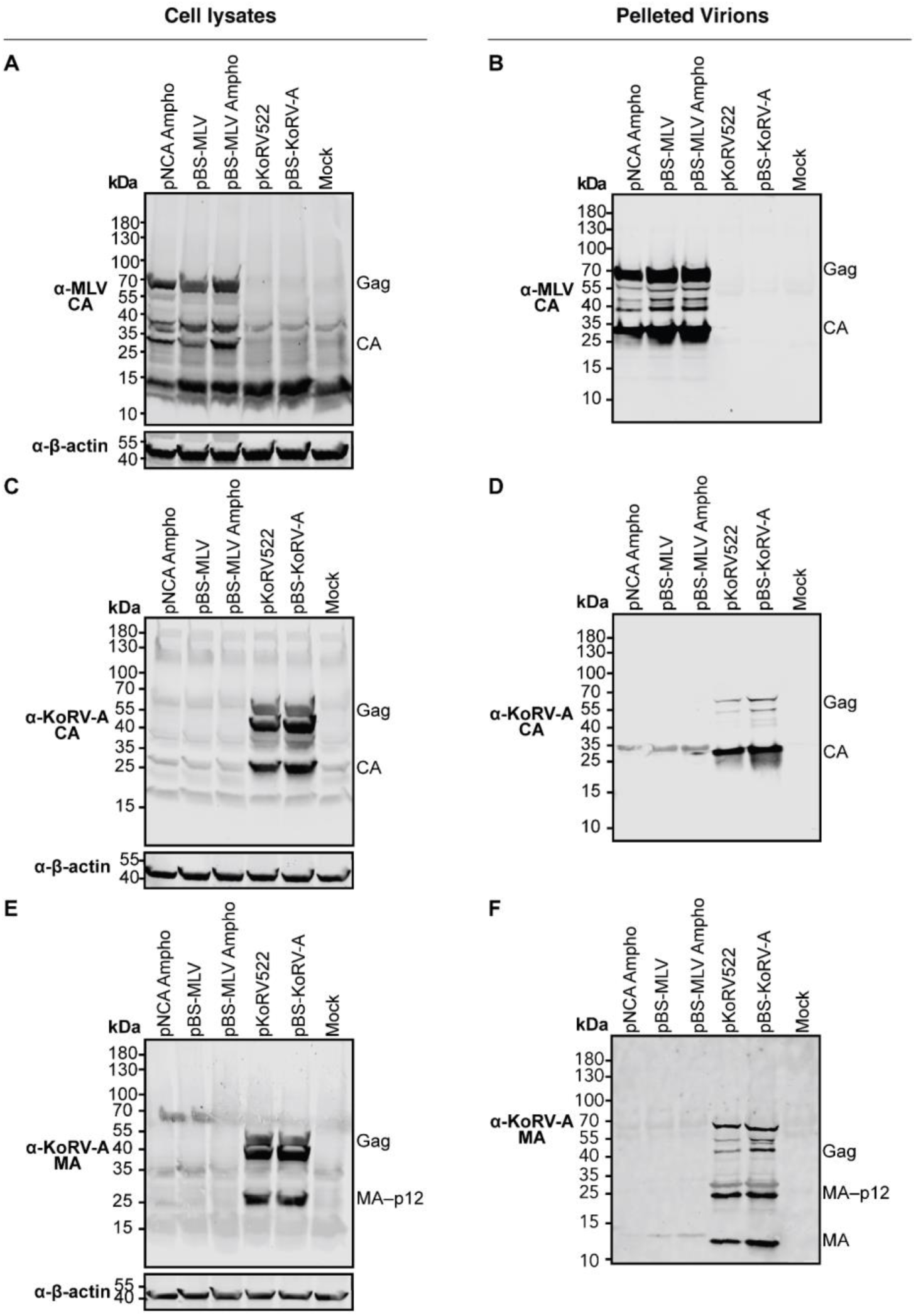
Expression and processing of KoRV-A and Mo-MLV Gag proteins in transfected cells and released virions. HEK293T cells were transfected with the indicated proviral plasmids. Cell lysates (A, C, E) and virions pelleted from equal volumes of culture supernatants by ultracentrifugation (B, D, F) were analyzed by Western blotting using antibodies against MLV CA (A, B), KoRV-A CA (C, D), or KoRV-A MA (E, F). Positions of the Gag precursor and processed CA (A–D), the MA-p12 processing intermediate (E, F), and mature MA (F) are indicated. β-Actin served as a loading control for cell lysates. Molecular masses are indicated in kDa.

Gammaretroviruses such as MLV utilize translational readthrough of the gag termination codon to generate the Gag–Pol fusion protein [54]. Western blot analysis using KoRV-A RT-and IN-specific antibodies detected the full-length KoRV-A Gag–Pol precursor in transfected cell lysates (Fig 4A and 4C; S6A, S6C, S6E, and S6G Fig). Mature KoRV-A RT (∼70 kDa) and IN (∼40 kDa) were detected in virions pelleted from culture supernatants (Fig 4B and 4D; S6B, S6D, S6F, and S6H Fig). Neither antibody cross-reacted with the corresponding Mo-MLV proteins. Together, these findings demonstrate that KoRV-A Gag–Pol is expressed and undergoes proteolytic processing. Having established that pBS-KoRV-A and the pBS-MLV constructs bearing ecotropic or amphotropic Env recapitulated the phenotypes of pKoRV522 and pNCA Ampho, respectively, these constructs were used in subsequent experiments. For simplicity, the corresponding viruses are hereafter referred to as KoRV-A and MLV unless otherwise specified.

**Fig 4.**
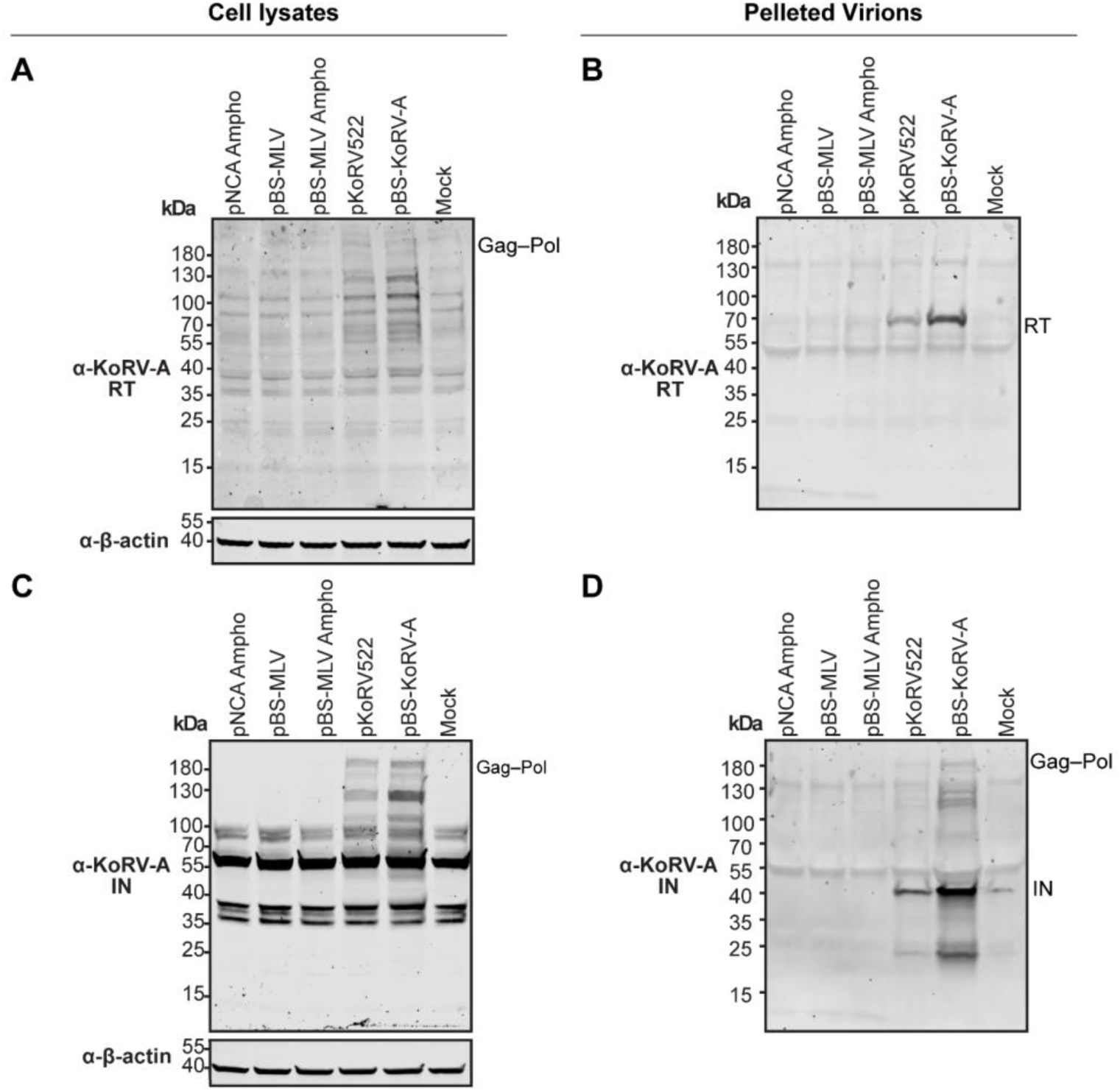
Detection of KoRV-A RT and IN in producer cell lysates and pelleted virions. (A, B) Western blot analysis of transfected cell lysates (A) and virions pelleted from equal volumes of culture supernatant (B) using a KoRV-A RT-specific antibody. (C, D) Western blot analysis of transfected cell lysates (C) and virions pelleted from equal volumes of culture supernatant (D) using a KoRV-A IN-specific antibody. β-Actin served as a loading control for cell lysates. Positions of the Gag–Pol precursor and mature RT and IN proteins are indicated. Molecular masses are indicated in kDa.

To ascertain whether there are differences in viral gene expression between KoRV-A and MLV, EGFP reporter constructs were generated by inserting EGFP either between the MA and p12 domains of Gag (MA-EGFP-p12) or between the RT and IN domains of Pol (RT-EGFP-IN) (Fig 5A). Insertion of EGFP at either position did not measurably alter RT activity of ecotropic MLV, and the replication and RT activity of amphotropic MLV (S7A, S7B, and S7D–G Fig). In contrast, KoRV-A MA-EGFP-p12 reduced RT activity relative to untagged KoRV-A (S7C Fig). This defect should therefore be considered when interpreting KoRV-A Gag-EGFP reporter data.

**Fig 5.**
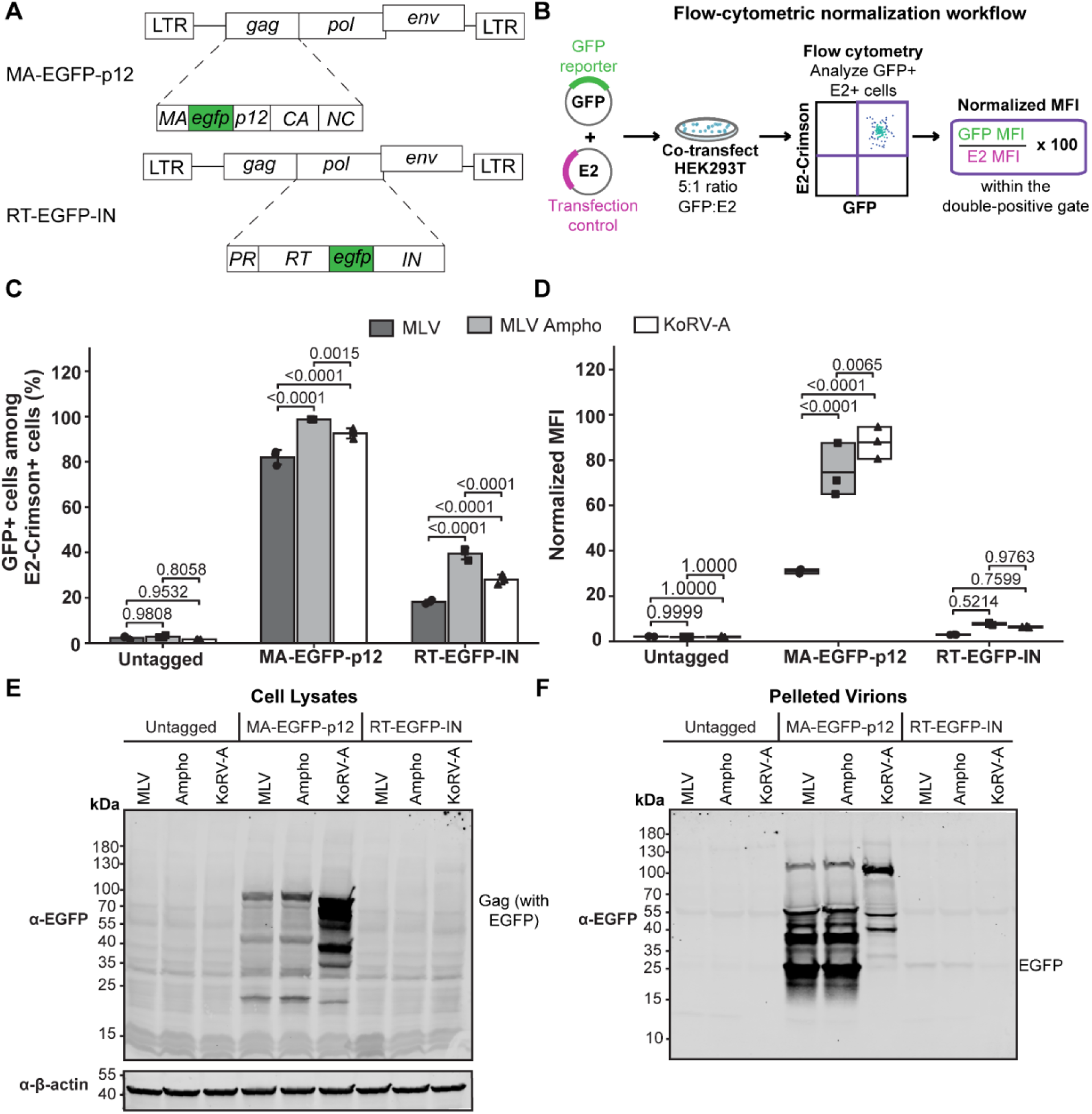
Assessment of KoRV-A and MLV Gag–Pol expression using EGFP reporter constructs. (A) Schematic representation of the EGFP-tagged proviral constructs. EGFP was inserted either between the MA and p12 domains of Gag or between the RT and IN domains of Pol. (B) Experimental design for flow-cytometric analysis. Cells were co-transfected with an EGFP-tagged proviral construct and an E2-Crimson expression plasmid at a 5:1 DNA ratio. Analyses were restricted to E2-Crimson-and EGFP-positive cells. Normalized MFI was calculated as 100 × the GFP median fluorescence intensity (MFI) divided by the E2-Crimson MFI within this double-positive population. (C) Percentage of EGFP-positive cells within the E2-Crimson-positive population following transfection with the untagged control or constructs containing EGFP between MA and p12 (MA–EGFP–p12) or between RT and IN (RT–EGFP– IN). Bars represent the mean ± SD. (D) Normalized GFP MFI within the E2-Crimson-and EGFP-positive population. Floating bars show the minimum and maximum values, horizontal lines indicate the medians. For (C) and (D), data were analyzed by ordinary two-way ANOVA, with virus and reporter construct as the two factors, followed by Šídák’s multiple-comparisons test comparing viruses within each construct. Exact Šídák-adjusted *p* values are shown. (E) Western blot analysis of transfected-cell lysates using an anti-EGFP antibody to assess intracellular expression of the EGFP-tagged proteins. β-Actin served as a loading control. (F) Western blot analysis of virions pelleted from equal volumes of culture supernatant using an anti-EGFP antibody to assess incorporation of the tagged proteins into released virions. Molecular masses are indicated in kDa.

Cells were co-transfected with an E2-Crimson expression plasmid to account for differences in transfection efficiency, and EGFP and E2-Crimson expression were quantified by flow cytometry (Fig 5B). The percentage of EGFP-positive cells among E2-Crimson-positive cells differed among the viruses and reporter constructs, although the direction and magnitude of these differences were not consistent across replicates (Fig 5C; S8A and S8C Fig). However, analysis of normalized EGFP median fluorescence intensity (MFI) in transfected cells consistently showed that KoRV-A exhibits greater cell-associated accumulation of EGFP-tagged proteins than MLV (Fig 5D; S8B and S8D Fig). This finding was supported by Western blot analysis of producer-cell lysates (Fig 5E; S9A and S9C Fig). KoRV-A RT-EGFP-IN also exhibited a modest increase in normalized MFI relative to the corresponding ecotropic MLV construct, although this difference was not statistically significant (Fig 5D; S8B and S8D Fig). The comparatively low abundance of the full-length Gag–Pol precursor, which is generated through translational readthrough, may limit the sensitivity of the RT-EGFP-IN reporter for detecting modest differences in Pol expression. The higher EGFP-positive percentage and normalized MFI observed for MLV Ampho in some experiments likely reflects contributions from secondary infection or viral spread in HEK293T cells, in addition to expression in initially transfected cells (Fig 5C–5D; S8A–D Fig). Western blot analysis of virions pelleted from equal volumes of cultured supernatant showed greater abundance of MA-EGFP-p12 derived products with ecotropic or amphotropic MLV compared to that of KoRV-A (Fig 5F; S9B and S9D Fig). Moreover, proteolytic processing of KoRV-A MA-EGFP-p12 was defective as the expected processed EGFP product was absent (Fig 5E-F; S9 Fig). This processing defect likely contributed to its reduced RT activity. With the RT-EGFP-IN constructs of KoRV-A and ecotropic and amphotropic MLV, no product was observed with the cell lysates likely due to reduced translation of the full-length Gag–Pol (Fig 5E; S9A and S9C Fig). However, in the pelleted supernatant, a band corresponding to EGFP is observed with the MLV constructs while this band is absent with KoRV-A, reflecting reduced extracellular accumulation of KoRV-A-derived proteins (Fig 5F; S9B and S9D Fig).

Because EGFP insertion altered RT activity in the KoRV-A Gag construct, these findings were independently confirmed using the smaller FLAG epitope tag. FLAG was inserted in either the p12 or RT in both KoRV-A and MLV, and results were consistent between the two insertion sites (S10A Fig). Unlike the KoRV-A MA-EGFP-p12 construct, insertion of the FLAG epitope at either site did not alter RT activity relative to untagged virus (S10B Fig). p12 FLAG-tagged constructs showed comparable expression of cell-associated Gag and Gag-derived proteolytic products between KoRV-A and MLV (S10C–E Fig). However, KoRV-A p12–FLAG tagged proteins accumulated less in the pelleted viral supernatant, mirroring the pattern observed with the EGFP reporters (Fig 5F; S9B and S9D Fig; S10F–H Fig). With the RT FLAG-tagged constructs, no protein products were detected in cell lysates, but a band corresponding to the mature RT was observed only in pelleted MLV virions, recapitulating the RT-EGFP-IN observations (Fig 5E–F; S9 Fig; S10C–H Fig). Catalytically inactive PR (D32A) mutants of the FLAG-tagged constructs were also analyzed to compare cell-associated and extracellular viral protein accumulation independent of proteolytic processing (S11 Fig). This allowed a direct comparison of KoRV-A and MLV Gag–Pol precursor accumulation without the complication of different rates of proteolytic activity. Western blot analyses of the p12–FLAG constructs of both retroviruses showed accumulation of the full-length Gag in the cell lysates (S11A–C Fig). In the supernatant, on the other hand, the Gag–Pol precursor and other protein products (likely proteolytic cleavage products derived from Gag and Gag–Pol precursor) were abundant while Gag was detected (S11D–F Fig). Although no products were observed in the cell lysates of the RT–FLAG constructs, the full-length Gag–Pol was detected in the supernatant with MLV but not with KoRV-A (S11 Fig). Together, these data demonstrate that the reduced replication phenotype of KoRV-A is not explained by diminished transcription or intracellular viral protein expression but instead reflects post-transcriptional differences affecting extracellular viral protein accumulation, a defect that persists even when proteolytic maturation is eliminated.

### Cooperation between PR and RT is required for efficient KoRV-A RT activity

A panel of KoRV-A/MLV chimeric viruses was generated in which individual or combined genomic regions were exchanged between the two viruses to identify viral elements responsible for the reduced exogenous RT activity of KoRV-A. Following transfection of HEK293T cells, particle-associated RT activity was quantified using an exogenous RT assay. Replacement of the entire *pol* region of KoRV-A with that of MLV increased RT activity by approximately 100-fold, whereas replacement of the complete *gag*–*pol*–*env* coding region restored RT activity to levels comparable to WT MLV (approximately 1,000-fold greater than WT KoRV-A) further illustrating that the LTR of KoRV-A can sufficiently support transcription (Fig 6A). Individual Pol domains were replaced to determine the minimal region needed for robust RT activity. Replacement of the KoRV-A PR domain, IN domain, or combined PR–IN region with the corresponding MLV sequences did not restore RT activity (Fig 6A). Notably, the IN-only and PR–IN chimeras exhibited approximately 100-fold and 10-fold lower RT activity than WT KoRV-A, respectively (Fig 6A). These findings demonstrate that the PR–RT domains must function together to restore RT activity.

**Fig 6.**
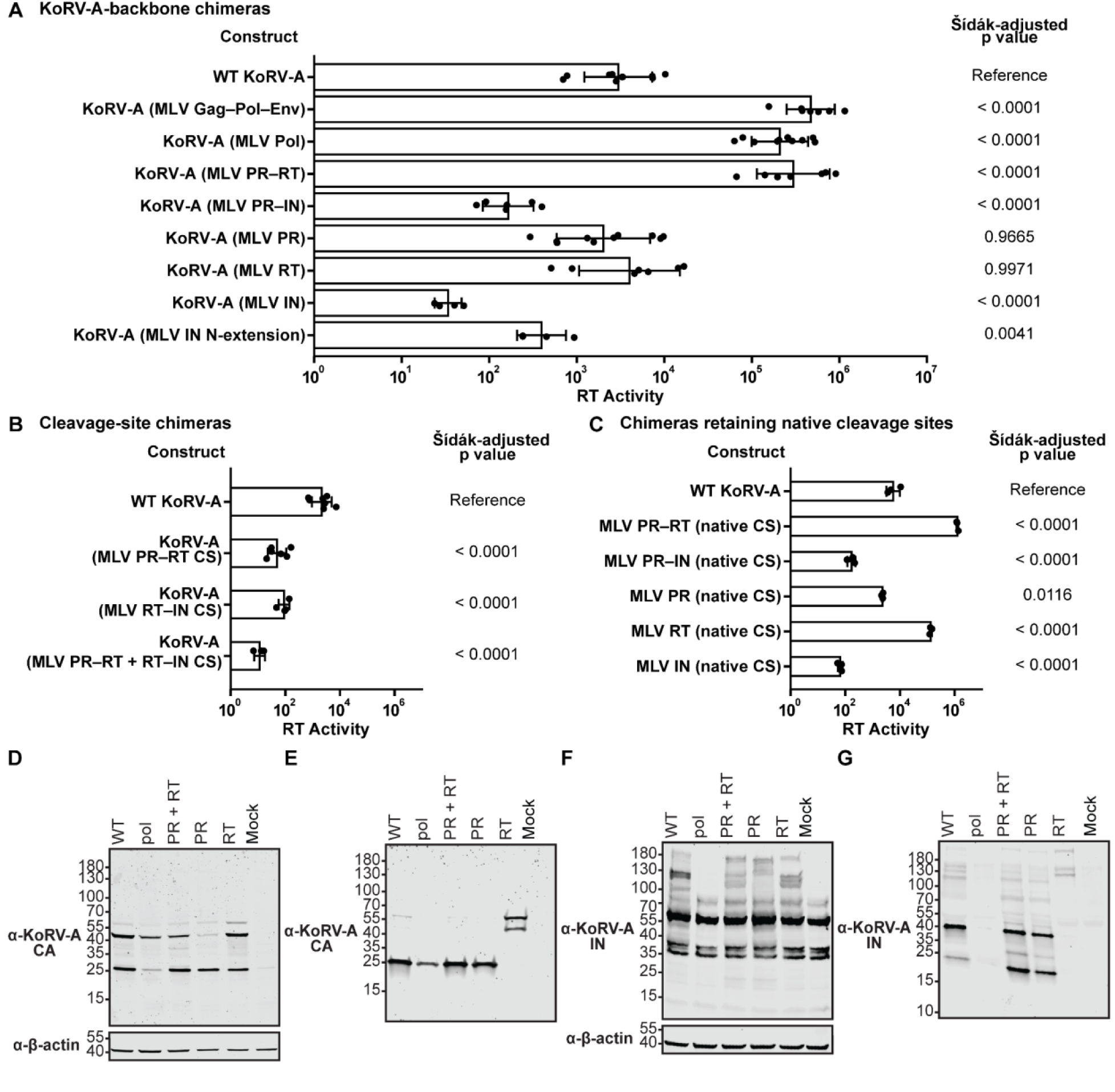
Characterization of KoRV-A chimeras containing MLV Pol domains and cleavage sites. (A) Exogenous RT activity in culture supernatants from HEK293T cells transfected with WT KoRV-A or the indicated KoRV-A-backbone chimeras containing MLV sequences. (B) Exogenous RT activity of KoRV-A chimeras containing MLV sequences at the PR–RT and/or RT–IN cleavage sites. (C) Exogenous RT activity of KoRV-A chimeras containing the indicated MLV Pol domains while retaining the native KoRV-A cleavage sites. RT activity is presented as the geometric mean ± geometric SD; individual points represent independent biological replicates. Statistical significance was assessed by ordinary one-way ANOVA followed by Šídák’s multiple-comparisons test comparing each chimera with WT KoRV-A. Exact Šídák-adjusted *p* values are shown. (D, E) Western blot analysis of matched transfected-cell lysates (D) and virions pelleted from equal volumes of culture supernatant (E) using an antibody against KoRV-A CA. (F, G) Western blot analysis of matched transfected-cell lysates (F) and pelleted virions (G) using an antibody against KoRV-A IN. β-Actin served as a loading control for cell lysates. Molecular masses are indicated in kDa.

Reciprocal chimeras generated in the MLV backbone revealed a distinct pattern. Replacement of the complete pol, the combined PR–RT region, the combined PR–IN region, or individual KoRV-A PR and RT domains into MLV all markedly reduced RT activity, indicating that KoRV-A Pol domains are broadly incompatible with the MLV replication machinery regardless of whether they are introduced individually or in combination (S12A Fig). Consistent with this context dependence, insertion of the MLV IN N-terminal extension into KoRV-A caused a modest but statistically significant reduction in RT activity (Fig 6A), whereas deletion of the corresponding N-terminal extension from MLV reduced RT activity by approximately 10-fold, although activity remained substantially higher than that of WT KoRV-A (S12A Fig). Thus, the MLV IN N-terminal extension contributes to RT activity in its native MLV context but is not sufficient to improve—and may modestly impair—RT activity when transferred into KoRV-A. Hence, the coordinated PR–RT pairing requirement observed in the KoRV-A backbone does not simply reverse when the direction of domain transfer is reversed, underscoring the context dependence of Pol domain function.

Because p12 is highly divergent between KoRV-A and MLV, particle-associated RT activity of reciprocal p12 chimeras was also examined to determine whether this Gag domain contributes to the difference in particle-associated RT activity. Substitution of KoRV-A p12 into MLV reduced RT activity by approximately 10-fold relative to WT MLV, although activity remained substantially higher than that of WT KoRV-A (S13B Fig). Conversely, substitution of MLV p12 into KoRV-A also reduced RT activity by approximately 10-fold rather than rescuing it (S13B Fig). Thus, exchanging p12 was detrimental in both viral backgrounds and did not account for the low RT activity of KoRV-A. These reciprocal effects further indicate that viral domains function within virus-specific structural and processing contexts.

### Cleavage-site sequence alone does not account for reduced KoRV-A RT activity

The sequences surrounding the PR–RT and RT–IN cleavage sites were exchanged between KoRV-A and MLV to investigate cleavage site compatibility. Substitution of either or both KoRV-A cleavage sites into MLV had no detectable effect on RT activity, demonstrating that these cleavage sequences are compatible with MLV PR (S13A Fig). Conversely, replacement of the corresponding KoRV-A cleavage sites with the corresponding MLV sequences significantly reduced KoRV-A RT activity (Fig 6B). Even when the native KoRV-A cleavage sites were retained, the PR-only, IN-only, and PR–IN chimeras all failed to rescue RT activity, with the IN-containing chimeras remaining particularly defective (Fig 6C). On the other hand, substitution of the KoRV-A RT with that of MLV while maintaining the native KoRV-A cleavage sites, increased RT activity by approximately 10-fold, but did not restore activity to the level of the complete Pol or PR–RT chimeras (Fig 6C). Thus, MLV RT confers greater particle-associated RT activity in the KoRV-A background, but does not fully explain the low RT activity of KoRV-A. The PR–RT chimera rescued RT activity by approximately 100-fold relative to WT KoRV-A regardless of whether KoRV-A or MLV cleavage sites were used, demonstrating that coordinated PR–RT function can overcome cleavage site constraints when both domains are optimized (Fig 6C). Thus, cleavage-site compatibility influences proteolytic processing and RT activity and efficient rescue requires coordinated interaction between PR and RT rather than exchange of either domain alone.

### Distinct regions of Pol independently regulate RT activity and particle-associated Gag/CA levels

Immunoblotting with an antibody against KoRV-A CA was used to assess whether restoration of RT activity was accompanied by changes in Gag expression and processing. Replacement of the complete KoRV-A Pol region with MLV Pol restored RT activity but markedly reduced cell-associated Gag and extracellular particle-associated CA (Fig 6D–E). WT KoRV-A and the PR-only and PR–RT chimeras exhibited comparable levels of cell-associated Gag and extracellular particle-associated CA, despite marked restoration of RT activity only in the PR–RT chimera (Fig 6D–E). Thus, the recovery of enzymatic activity was not attributable to increased steady-state levels of particle-associated Gag or CA. However, immunoblotting has a limited dynamic range; thus, more modest differences in viral protein production or particle release among the constructs cannot be excluded. By contrast, the RT-only chimera exhibited defective proteolytic processing, illustrating incompatibility between the exchanged MLV domain, its associated cleavage sites, and the KoRV-A PR (Fig 6D–E). This interpretation is consistent with the cleavage-site exchange experiments, which demonstrated that KoRV-A does not efficiently tolerate MLV cleavage-site sequences. Cell lysates and pelleted virions of the MLV backbone chimeras were similarly assessed with a Western blot. Exchange of MLV Pol with KoRV-A Pol reduced both cell-associated and extracellular viral protein levels (S12B–G Fig). MLV chimeras containing KoRV-A PR or KoRV-A PR–RT exhibited defective proteolytic processing, further demonstrating that KoRV-A PR cannot efficiently process substrates containing MLV cleavage-site sequences. In contrast, insertion of KoRV-A RT into the MLV backbone did not disrupt proteolytic processing. Overall, these results further implicate PR–RT compatibility and proteolytic processing as major determinants of the RT activity of released particles.

The pattern of IN proteolytic processing across the chimera panel was investigated by immunoblotting of transfected-cell lysates and pelleted virions using the KoRV-A IN-specific antibody. In cell lysates, the full-length Gag–Pol precursor and several smaller products were detected for WT KoRV-A and most chimeras, whereas some processed species were absent from the RT-only chimera, consistent with broader disruption of Gag–Pol processing in this construct (Fig 6F). In pelleted virions, an approximately 25-kDa IN-reactive product was detected for WT KoRV-A and the PR–RT and PR-only chimeras (Fig 6G). No specific IN signal was detected for the complete MLV Pol chimera, consistent with the lack of cross-reactivity of the KoRV-A IN-specific antibody with MLV IN rather than a biological difference. Neither mature IN nor the approximately 25-kDa product was detected for the RT-only chimera (Fig 6G).

Immunoblotting of the catalytically inactive PR (D32A) mutant further supported the dependence of these processing events on viral PR activity. Using the KoRV-A CA-specific antibody, full-length Gag accumulated in both cell lysates and pelleted virions from the PR mutant, whereas processed CA was readily detected for WT KoRV-A but not the mutant (S14A and S14C Fig). Similarly, the full-length Gag–Pol precursor was more distinct in cell lysates from the PR mutant than in WT KoRV-A when probed with the KoRV-A IN-specific antibody, consistent with accumulation of the uncleaved precursor when PR activity was abolished (S14B Fig). The approximately 25-kDa IN-reactive species was also absent from the PR mutant (S14B and S14D Fig), supporting its assignment as a PR-dependent processing product rather than a nonspecific degradation product. Although the identity and biological function of this species remain unknown, these observations suggest that KoRV-A IN may undergo additional proteolytic processing beyond that previously recognized.

### Extracellular virion density and spatial organization differ between KoRV-A, MLV, and Pol-domain chimeras

Cells transfected with MLV, KoRV-A, or the KoRV-A Pol-domain chimeras (containing the corresponding MLV cleavage site sequences flanking the exchanged regions) were examined by TEM. Compared with MLV, KoRV-A displayed a visually apparent increase in tightly clustered extracellular virions, with multiple particles frequently observed immediately adjacent to one another (Fig 7A). Although the PR–RT chimera restored RT activity, virion aggregation persisted with larger clusters as extracellular virions were more abundant. The PR-only chimera closely resembled WT KoRV-A in both particle abundance and morphology. The RT-only chimera also exhibited increased extracellular virion abundance relative to WT KoRV-A; however, most, if not all, particles in sampled TEM fields displayed an immature morphology characterized by the absence of condensed electron-dense cores (Fig 7A). In contrast, relatively few virions were observed in cells transfected with the KoRV-A p12 chimera. The particles that were present displayed mature morphology without obvious membrane-associated stalks or tethered budding profiles, providing no morphological evidence of a classical L-domain-dependent budding arrest [55–58] (S13C Fig). However, TEM observations alone cannot exclude a more subtle defect in particle assembly or release.

**Fig 7.**
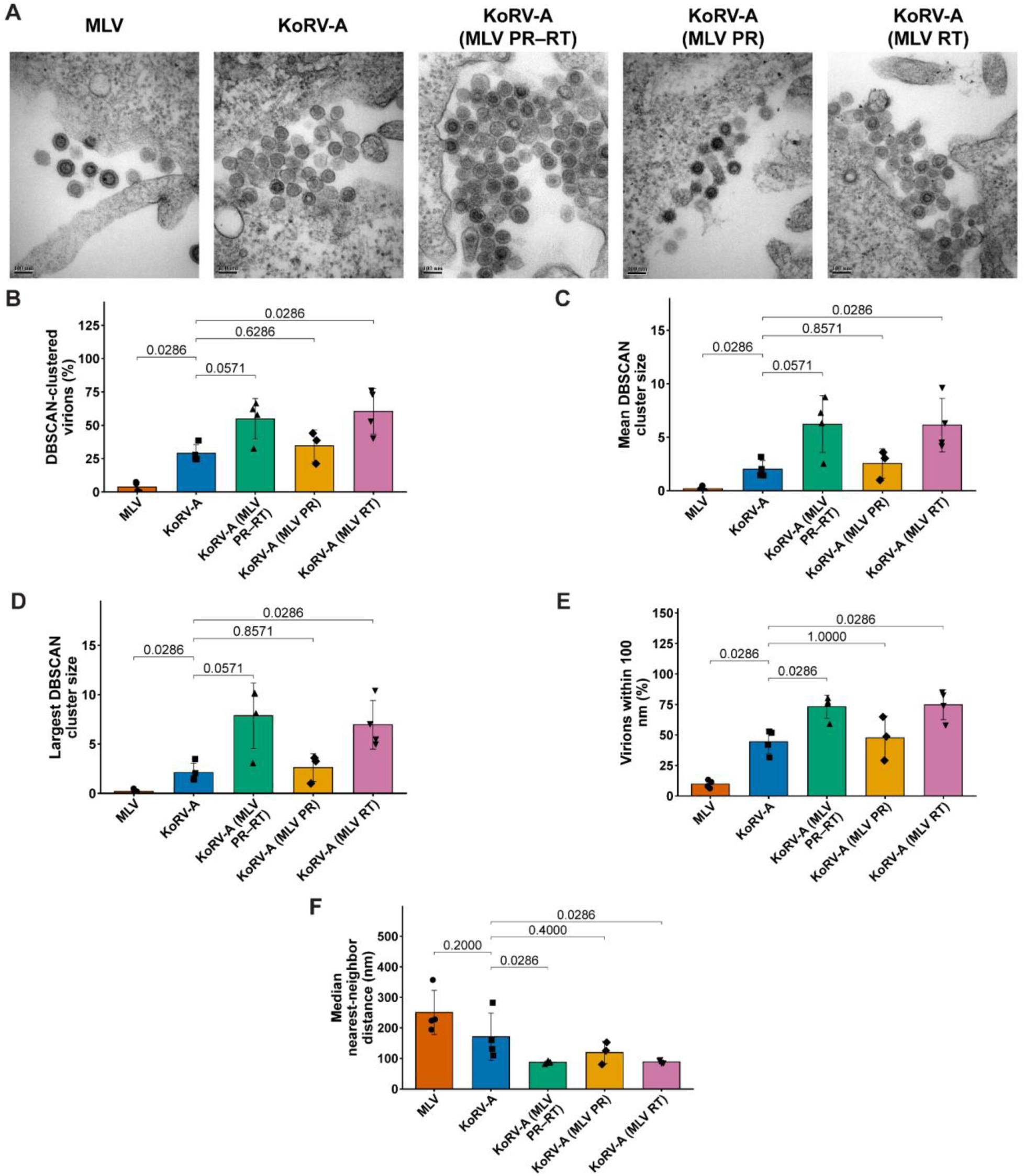
Virion clustering of MLV, KoRV-A, and KoRV-A constructs containing MLV PR and/or RT. (A) Representative TEMs of MLV, KoRV-A, KoRV-A with MLV PR–RT, KoRV-A with MLV PR, and KoRV-A with MLV RT, shown from left to right. Scale bars, 100 nm. (B) Percentage of virions assigned to a DBSCAN cluster. (C) Mean DBSCAN cluster size. (D) Largest DBSCAN cluster size. (E) Percentage of virions with a nearest neighbor within 100 nm. (F) Median nearest-neighbor distance. The numbers of biological replicates and analyzed TEM fields, respectively, were: MLV, 4 and 129; KoRV-A, 4 and 86; KoRV-A (MLV PR–RT), 4 and 99; KoRV-A (MLV PR), 3 and 67; and KoRV-A (MLV RT), 4 and 72. Bars represent the mean ± SD, and individual symbols represent biological replicates. Statistical comparisons were performed on biological-replicate summaries using two-sided Mann–Whitney U tests. *P* values are unadjusted and are presented as exploratory analyses of Pol-component effects on virion organization. These biological-replicate-level comparisons are complemented by exploratory image-level regression analyses adjusted for virion abundance (S18 Fig).

Replacement of the complete KoRV-A Pol region with MLV Pol, as well as the PR–IN and IN-only substitutions, markedly reduced extracellular virion abundance. Quantitative spatial analysis was therefore limited to TEM images of cells transfected with MLV, KoRV-A, or the KoRV-A chimeras containing MLV RT, PR, or PR–RT. Images containing a single virion were retained for particle-abundance and DBSCAN analyses, although nearest-neighbor distances could not be calculated for these images. The numbers of biological replicates, TEM images, and virions analyzed are summarized in S3 Table. For each image, we calculated the percentage of DBSCAN-clustered virions, number of clusters, mean and maximum cluster sizes, percentage of virions with a nearest neighbor within 100 nm, and median nearest-neighbor distance.

Supplementary figures provided detailed views of virion abundance and spatial organization. Replicate-level distributions of mean virions per image are shown in S15A Fig, individual virion nearest-neighbor distances in S15B Fig (presented descriptively because of non-independence within replicates and images), and virion counts per image in S15C Fig. Image-level relationships between virion abundance and spatial measurements are shown in S15D–S15H Fig. Substantial image-to-image heterogeneity was evident across all experimental groups, with individual TEM fields ranging from no detectable clustering to highly clustered arrays. Despite this heterogeneity, KoRV-A and the KoRV-A chimeras showed a consistent pattern: more imaging fields with elevated DBSCAN-clustered particles, larger mean and maximum cluster sizes, and greater proportions of particles within 100 nm of another virion compared to MLV. This pattern was most pronounced in the PR–RT and RT-only chimeras, which exhibited both the highest virion abundance per image and the highest cluster-based measurements, along with the shortest median nearest-neighbor distances.

Pairwise comparisons of biological-replicate summaries were performed using two-sided Mann– Whitney U tests at the replicate level. Because these comparisons were exploratory, *p* values were not adjusted for multiple comparisons; exact unadjusted p values are shown in Fig 7 and should be interpreted alongside the direction and magnitude of the observed differences.

WT KoRV-A exhibited substantially greater clustering than MLV across multiple metrics. Compared to MLV, KoRV-A had a higher percentage of DBSCAN-clustered virions (29.0% versus 3.7%; Fig 7B), larger mean cluster size (2.01 versus 0.20 virions; Fig 7C), and larger maximum cluster size (2.11 versus 0.20 virions; Fig 7D). A greater proportion of KoRV-A virions also had a neighboring particle within 100 nm (44.5% versus 9.8%; Fig 7E). Although the mean replicate - level median nearest-neighbor distance was numerically lower for KoRV-A (170.8 versus 250.7 nm; Fig 7F), the exploratory comparison did not provide evidence of a statistically significant difference. KoRV-A additionally exhibited more clusters per image (0.46 versus 0.05; S3 Table) and more virions per image (5.4 versus 3.4; S15C Fig).

The PR–RT chimera showed substantially greater particle abundance and clustering than WT KoRV-A. Compared to KoRV-A, the PR–RT chimera had a higher percentage of DBSCAN-clustered particles (54.7% versus 29.0%; Fig 7B), larger mean and maximum cluster sizes (6.22 versus 2.01 virions and 7.86 versus 2.11 virions, respectively; Fig 7C–D), a greater proportion of particles within 100 nm (73.2% versus 44.5%; Fig 7E), and a shorter mean replicate-level median nearest-neighbor distance (87.2 versus 170.8 nm; Fig 7F). In contrast, the PR-only chimera did not substantially alter spatial organization compared to WT KoRV-A. The two groups showed minimal differences in percentage of DBSCAN-clustered particles (34.6% versus 29.0%; Fig 7B), mean cluster size (2.54 versus 2.01 virions; Fig 7C), maximum cluster size (2.62 versus 2.11 virions; Fig 7D), proportion within 100 nm (47.6% versus 44.5%; Fig 7E), and mean replicate-level median nearest-neighbor distance (119.5 versus 170.8 nm; Fig 7F). Cluster density per image was also similar (0.53 versus 0.46; S3 Table), as was virion abundance (6.1 versus 5.4; S15C Fig). The RT-only chimera, like the PR–RT chimera, displayed substantially greater particle abundance and clustering than WT KoRV-A. Compared to KoRV-A, the RT-only chimera had higher particle abundance (12.4 versus 5.4 virions per image; S15C Fig), more clusters per image (1.27 versus 0.46; S3 Table), a higher percentage of DBSCAN-clustered particles (60.5% versus 29.0%; Fig 7B), larger mean and maximum cluster sizes (6.13 versus 2.01 virions and 6.95 versus 2.11 virions, respectively; Fig 7C–D), a greater proportion of particles within 100 nm (74.8% versus 44.5%; Fig 7E), and a shorter mean replicate-level median nearest-neighbor distance (88.2 versus 170.8 nm; Fig 7F). Together, these findings demonstrate that virion aggregation is not abolished when KoRV-A Pol domains are replaced with MLV sequences. Instead, the RT and PR–RT domains substantially enhance the clustering phenotype, whereas the PR domain alone does not drive spatial reorganization.

For visualization, S16 pools particle groups across TEM images and biological replicates, whereas S17 pools virion-level nearest-neighbor distances. Single particle groups were retained in S16 Fig. In contrast, images containing only one virion were excluded from S17 Fig because a nearest-neighbor distance could not be calculated.

Analysis of DBSCAN particle-group-size distributions revelated striking differences in how virions organize extracellularly. MLV showed a skewed distribution toward single virions: 93.3% of particle groups consisted of single virions, 5.6% of pairs, and only 1.1% of larger groups (S16A Fig). In contrast, KoRV-A exhibited a broader distribution with 72.8% singles, 11.8% pairs, and 15.4% containing three or more virions, including groups exceeding 11 particles (S16B Fig). This shift toward larger assemblies persisted across the KoRV-A chimeras, though with varying intensity. The PR-only chimera most closely resembled WT KoRV-A (70.1% singles, 12.4% pairs, 17.5%≥3; S16D Fig), whereas the PR–RT and RT-only chimeras showed the greatest enrichment of larger groups (60.6% and 51.6^ singles, 12.4% pairs and 15.9% pairs, 24.9% and 32.5%≥3; S16C and S16E Fig). Exploratory comparisons of pooled, unbinned particle-group sizes confirmed differences between WT KoRV-A and MLV (Holm-adjusted p < 0.0001), the PR–RT chimera (p = 0.0005), and the RT-obnly chimera (p < 0.0001), but not the PR-only chimera (p =0.4330). These distributions consistently demonstrated larger extracellular particle assemblies for KoRV-A than MLV, with the most pronounced shift toward larger groups observed in the PR– RT and RT-only chimeras.

Nearest-neighbor distance distributions also differed among the groups. MLV virions were predominantly dispersed, with the largest proportion (44.4%) having nearest neighbors at 100 – 150 nm distance and only 16.3% within 100 nm (S17A Fig). KoRV-A showed dramatically different spacing: the predominant interval was 50–100 nm (58.5% of virions), with 61.1% total within 100 nm—nearly fourfold the proportion observed for MLV (S17B Fig). The PR-only chimera exhibited similarly close spacing to KoRV-A (57.3% in the 50–100 nm interval; S17D Fig), whereas the PR– RT and RT-only chimeras showed even greater concentration of short distances (77.2% and 82.7%, respectively; S17C and S17E Fig). Long-range distances (>300 nm) showed the inverse pattern: MLV had substantially more (19.2%) compared to KoRV-A (8.1%), the PR-only chimera (7.5%), and the PR–RT and RT-only chimeras (1.1% and 1.5%, respectively). Exploratory comparisons of pooled, unbinned nearest-neighbor distances confirmed differences between WT KoRV-A and MLV (Holm-adjusted p < 0.0001), and both PR–RT and RT-only chimeras (p < 0.0001 each), and the PR-only chimera (p = 0.0112). These distributions consistently demonstrated closer extracellular particle spacing for KoRV-A and its chimeras compared to MLV, with the most dramatic shift toward close proximity observed in the PR–RT and RT-only chimeras. Adjustment for particle abundance substantially attenuated the differences between WT KoRV-A and the chimeric constructs. For both the PR–RT and RT-only chimeras, density-adjusted comparisons showed no significant differences in the percentage of DBSCAN-clustered particles, mean cluster size, maximum cluster size, or median nearest-neighbor distance (S18A–C, E Fig). The sole exception was the proportion of particles with a nearest neighbor within 100 nm, which remained significantly different for the PR–RT chimera (S18D Fig). In contrast, the PR-only chimera showed no significant density-adjusted differences from WT KoRV-A across any spatial metric (S18A–E Fig).

In contrast to the chimeras, the density-adjusted clustering differences between WT KoRV-A and MLV persisted across multiple measurements. KoRV-A retained a greater percentage of DBSCAN-clustered particles, larger mean cluster sizes, and a greater proportion of particles within 100 nm, whereas median nearest-neighbor distance did not differ significantly. These findings indicate that WT KoRV-A possesses an intrinsic spatial organization phenotype distinct from MLV that is not explained by virion abundance alone.

Density adjustment affected the comparisons differently but did not establish distinct underlying mechanisms. For the PR–RT and RT-only chimeras, attenuation after adjustment indicates that their spatial measurements were closely associated with the greater numbers of virions detected per image. This analysis cannot distinguish whether particle abundance acts as a confounder, a mediator of the construct-associated phenotype, or a feature influenced by image sampling. In contrast, several differences between WT KoRV-A and MLV remained after adjustment for measured particle counts, indicating that these associations were not fully explained by the number of virions detected per image. Nevertheless, the adjusted results do not establish an intrinsic difference in virion organization or identify the biological processes responsible for the remaining associations.

Based on these observations, we propose a model in which MLV particles are efficiently released and dispersed into the extracellular environment, whereas KoRV-A particles remain concentrated near producer cells (Fig 8). Introduction of the MLV PR–RT region into KoRV-A increases extracellular particle abundance near cells but does not confer MLV-like dispersal, suggesting increased assembly or budding combined with persistent retention or inefficient dispersal. Because TEM quantifies particles within sampled imaging fields rather than total supernatant release, we cannot definitively distinguish among altered assembly, budding, release, retention, and dispersal mechanisms. These possibilities will require direct measurement of cell-associated versus freely dispersed particle populations.

**Fig 8.**
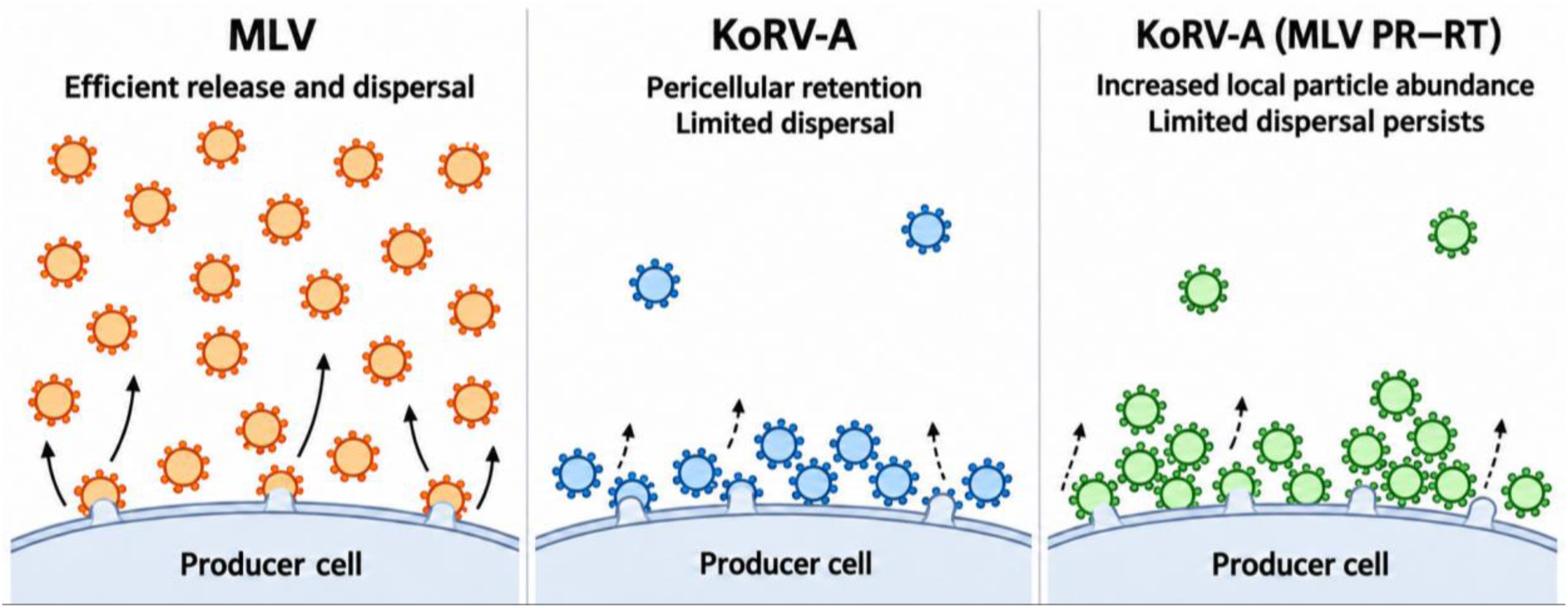
Proposed model for differences in extracellular particle organization of MLV, KoRV-A, and the KoRV-A PR–RT chimera. MLV particles appeared relatively dispersed in sampled extracellular fields. KoRV-A particles accumulated near producer cells, which could reflect reduced dispersal, increased pericellular retention, or another unmeasured process. The KoRV-A PR–RT chimera exhibited greater local particle abundance than WT KoRV-A while retaining a pericellular distribution. The experiments did not distinguish among altered particle production, release, dispersal, or retention.

## Discussion

KoRV-A is undergoing endogenization in koalas and is associated with diseases that threaten this endangered species, yet its replication biology remains poorly understood. Here, we investigated the mechanisms underlying its inefficient replication in laboratory experiments using human cell lines. By comparing KoRV-A with MLV and analyzing a series of Pol-modified viruses, we identified multiple genetically separable post-transcriptional phenotypes. KoRV-A exhibited markedly reduced particle-associated RT activity and viral spread, as well as differences in particle abundance and extracellular organization. These findings indicate that its limited replication cannot be attributed to a single impaired step.

The Pol-chimera experiments demonstrate that particle-associated RT activity depends on coordinated interactions among PR, RT, and their cleavage sites. Replacement of the combined PR–RT region with the corresponding MLV sequence increased particle-associated RT activity by approximately 100-fold. Replacing KoRV-A RT with MLV RT while retaining the native KoRV-A cleavage sites produced a more modest, approximately 10-fold increase. Thus, the catalytic properties of KoRV-A RT may contribute to its low activity but do not fully explain the defect. The greater rescue achieved by replacing both PR and RT suggests additional contributions from proteolytic processing, interactions between these domains, or Pol precursor folding.

Introducing MLV RT with its flanking MLV cleavage-site sequences disrupted proteolytic processing and virion maturation, consistent with inefficient processing of these heterologous sites by KoRV-A PR. This incompatibility was asymmetric: MLV PR tolerated the corresponding KoRV-A cleavage sites, whereas KoRV-A PR did not tolerate the MLV sequences. Introduction of KoRV-A Pol domains into MLV also broadly reduced RT activity, and transfer of the MLV IN N-terminal extension into KoRV-A was insufficient to improve activity. These findings support a model in which KoRV-A RT activity depends on coadapted interactions among Pol domains and cleavage sites.

Pol substitutions had distinct effects on RT activity, virion maturation, and particle abundance, indicating that these phenotypes are genetically separable. The PR–RT chimera restored RT activity and had a predominantly mature virion morphology. The RT-only chimera also increased extracellular virion counts by TEM but the particles remained defective in processing, maturation, and RT activity. Replacement of the complete KoRV-A Pol region restored RT activity while reducing intracellular Gag, extracellular CA, and extracellular virion abundance. Thus, increases in RT activity did not consistently accompany increases in Gag abundance or the number of extracellular particles.

TEM analysis also revealed differences between KoRV-A and MLV in extracellular particle organization. KoRV-A exhibited a greater proportion of particles within 100 nm of another particle, a higher percentage of DBSCAN-clustered particles, and larger mean and maximum cluster sizes. Differences in the proportion of particles within 100 nm, the percentage of DBSCAN-clustered particles, and mean cluster size persisted after adjustment for particle abundance. These findings suggest that the observed differences in clustering are not explained entirely by the number of virions detected per image.

The RT-only and PR–RT chimeras exhibited greater local extracellular virion abundance and apparent clustering than WT KoRV-A, whereas the PR-only chimera broadly resembled KoRV-A. The replicate-level comparisons estimate the overall extracellular phenotype associated with each construct, encompassing differences in both particle abundance and spatial organization. By contrast, the density-adjusted models estimate differences in spatial measurements among images with similar virion counts. Adjustment attenuated most differences involving the RT-only and PR–RT chimeras, indicating that their apparent clustering was closely associated with increased local particle abundance. However, virion abundance may be a mediator of the genotype-associated phenotype rather than simply a confounder. These analyses cannot determine whether the exchanged Pol domains affect particle production, assembly, release, dispersal, or retention near producer cells, nor whether increased proximity arises primarily as a geometric consequence of greater particle abundance. The unadjusted and density-adjusted analyses should therefore be interpreted together as evidence of an abundance-associated spatial phenotype, with limited evidence that the chimeras independently alter virion organization after accounting for particle number (Fig 8).

Several limitations should be considered. MLV is a laboratory-adapted virus and is therefore an imperfect comparator for a retrovirus undergoing endogenization. The experiments were conducted in human cell lines, and KoRV-A may behave differently in koala-derived cells. Particle-associated RT activity is a composite measurement that can reflect particle abundance, RT incorporation and processing, and enzymatic activity; it does not directly measure the intrinsic catalytic activity of KoRV-A RT. Similarly, TEM particle counts describe virions within selected imaging fields and do not directly measure total particle production or release. Although adjustment for particle abundance helps distinguish its contribution to spatial measurements, it does not identify the mechanisms underlying particle accumulation or clustering. Finally, although the low-RT-activity phenotype was confirmed in two independent KoRV-A clones, the mechanistic experiments employed the consensus KoRV-A molecular clone.

Together, these findings identify multiple genetically distinguishable limitations on KoRV-A replication. Compatibility among PR, RT, and the cleavage sites surrounding RT contributes to particle-associated RT activity, proteolytic processing and virion maturation. KoRV-A also exhibited a distinctive extracellular organization, with particles occurring in larger and denser groups than MLV particles. The persistence of clustering among the Pol-domain chimeras suggests that the exchanged Pol sequences alone do not account for this phenotype and implicates other viral or cellular factors. Env is a plausible contributor because it is displayed on the virion surface, but the present experiments do not establish that it mediates aggregation. Future studies could test whether KoRV-A Env is sufficient to alter extracellular particle organization and whether it is necessary for virion aggregation. These results provide a framework for understanding the limited replication of KoRV-A while identifying the determinants and consequences of extracellular particle clustering as important unresolved questions.

## Acknowledgements

We are grateful to Takayuki Nitta for providing the pKoRV522 plasmid. We thank Heinrich Göttlinger, Athma Pai, Craig Peterson, William Theurkauf, Leonid Yurkovetskiy, and Phillip Zamore for feedback and assistance with experimental design. We also thank members of the Luban laboratory, including Jida Huang, Javier Jaimes, Babatunde Olusola, and Karthika Nagalekshmi, for helpful discussions and assistance with experiments. We especially thank Keith Reddig and KyoungHwan Lee of the UMass Chan Medical School Electron Microscopy Core for assistance with transmission electron microscopy and acknowledge the UMass Chan Medical School Flow Cytometry Core for assistance with flow cytometry. Instrumentation used for TEM was supported by NIH Shared Instrumentation Grant S10OD025113. This work was supported by a Mello Fellowship awarded to M.A. and NIH grants R37AI147868 and U54AI170856 awarded to J.L.

## Data Availability Statement

All data underlying the findings reported in this study are provided within the manuscript, its Supporting Information files, or the accompanying Dryad Digital Repository deposit under the reserved DOI https://doi.org/10.5061/dryad.cjsxksnpj. The deposited materials include source data underlying the flow cytometry data, biological-replicate measurements, TEM images, TEM-derived particle coordinates and image-level spatial measurements, analysis scripts, and complete plasmid sequences.

---

**S1 Fig.**
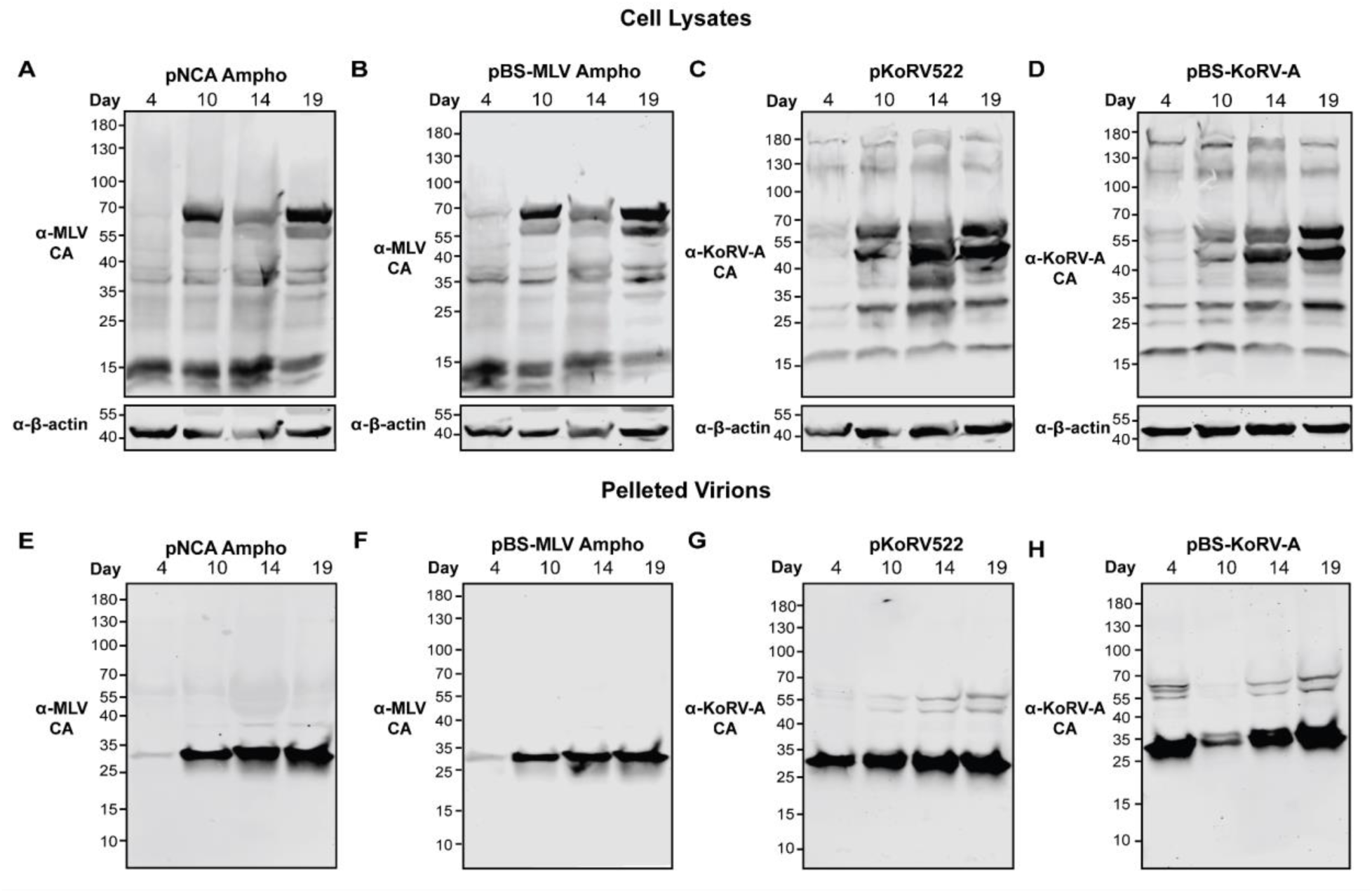
Western blot analysis of infected HEK293T cells and pelleted virions collected at 4, 10, 14, and 19 days post-infection. (A–D) Cell lysates from cultures infected with virus generated from pNCA Ampho (A), pBS-MLV Ampho (B), pKoRV522 (C), or pBS-KoRV-A (D). (E–H) Virions pelleted from equal volumes of the corresponding culture supernatants from cells infected with virus generated from pNCA Ampho (E), pBS-MLV Ampho (F), pKoRV522 (G), or pBS-KoRV-A (H). Samples were analyzed by Western blotting using antibodies against MLV CA or KoRV-A CA, as appropriate. β-Actin served as a loading control for cell lysates. Molecular masses are indicated in kDa.

**S2 Fig.**
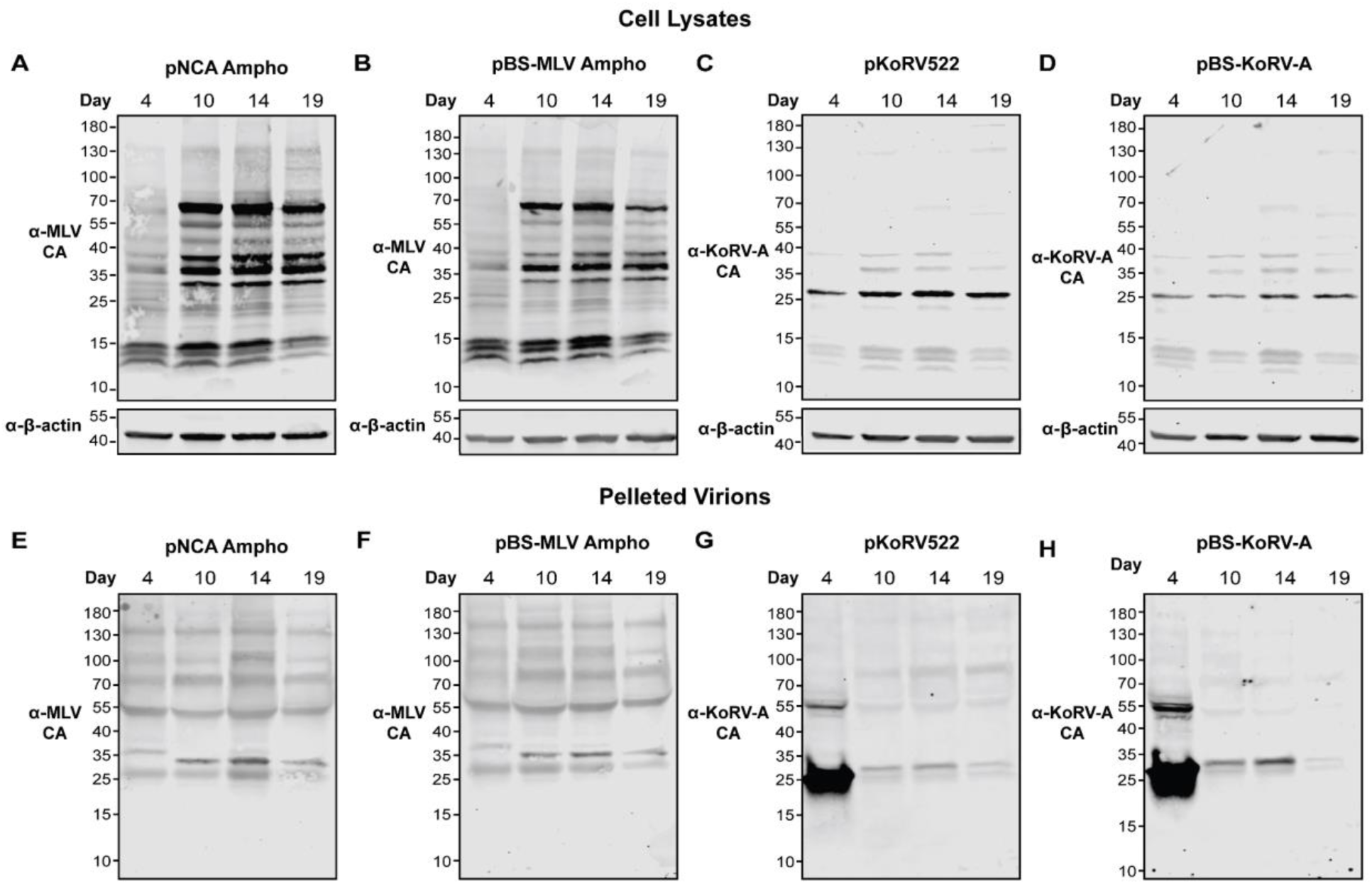
Western blot analysis of infected Jurkat cells and pelleted virions collected at 4, 10, 14, and 19 days post-infection. (A–D) Cell lysates from cultures infected with virus generated from pNCA Ampho (A), pBS-MLV Ampho (B), pKoRV522 (C), or pBS-KoRV-A (D). (E–H) Virions pelleted from equal volumes of the corresponding culture supernatants from cells infected with virus generated from pNCA Ampho (E), pBS-MLV Ampho (F), pKoRV522 (G), or pBS-KoRV-A (H). Samples were analyzed by Western blotting using antibodies against MLV CA or KoRV-A CA, as appropriate. β-Actin served as a loading control for cell lysates. Molecular masses are indicated in kDa.

**S3 Fig.**
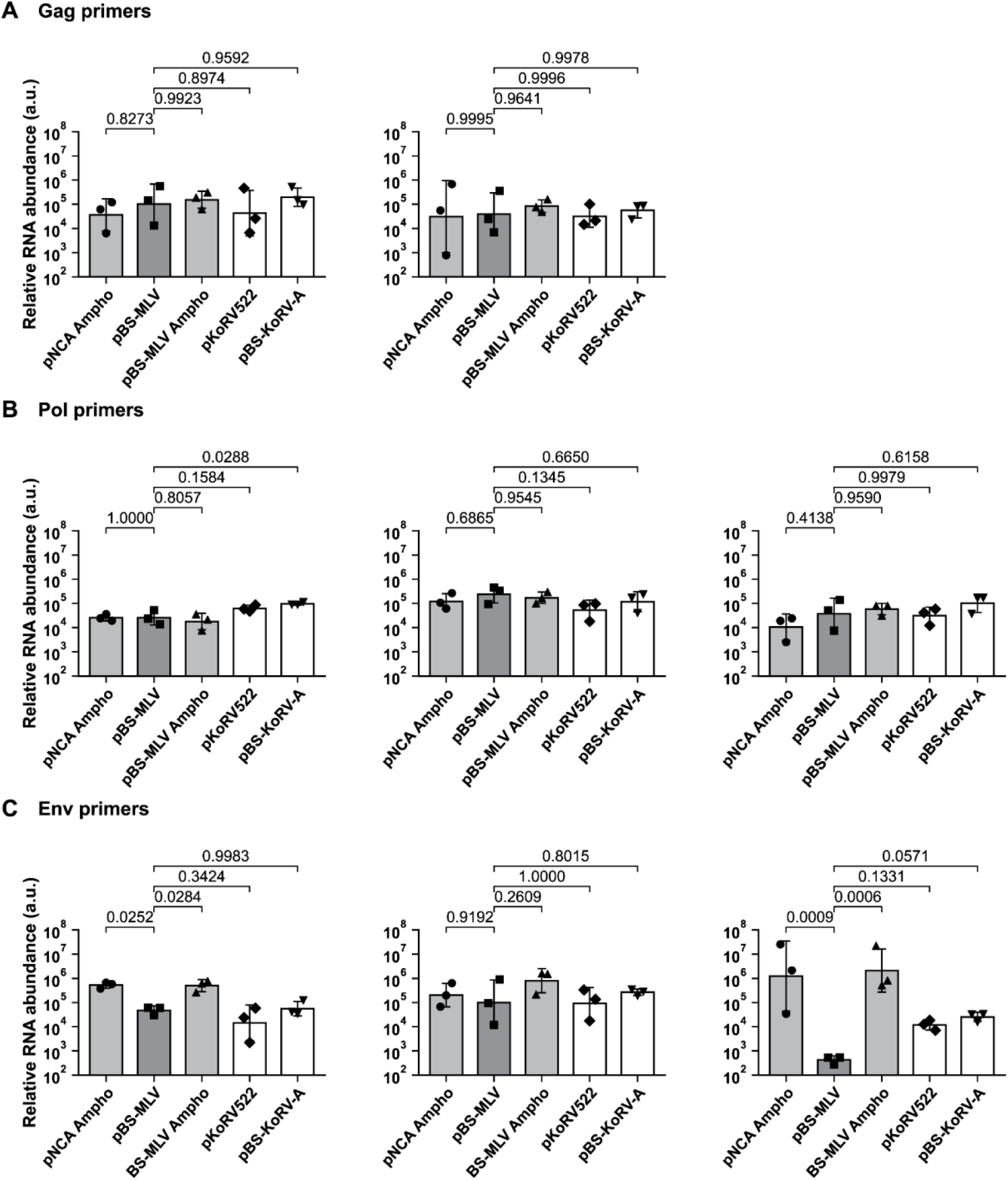
Quantification of KoRV-A and MLV viral transcript abundance by RT-qPCR. (A–H) Relative abundance of *gag* (A, B), *pol* (C–E), and *env* (F–H) transcripts in HEK293T cells transfected with the indicated KoRV-A or MLV proviral plasmids. Total RNA was isolated 72 h post-transfection, reverse transcribed, and analyzed by quantitative PCR using gene-specific primer pairs. Transcript abundance was normalized to β-actin and calculated relative to the mock-transfected control. Bars represent geometric means ± geometric standard deviations, and individual measurements are overlaid. Statistical significance was evaluated separately for each panel using a lognormal one-way ANOVA followed by Dunnett’s multiple-comparisons test comparing each construct with pBS-MLV. Exact Dunnett-adjusted *p* values are shown.

**S4 Fig.**
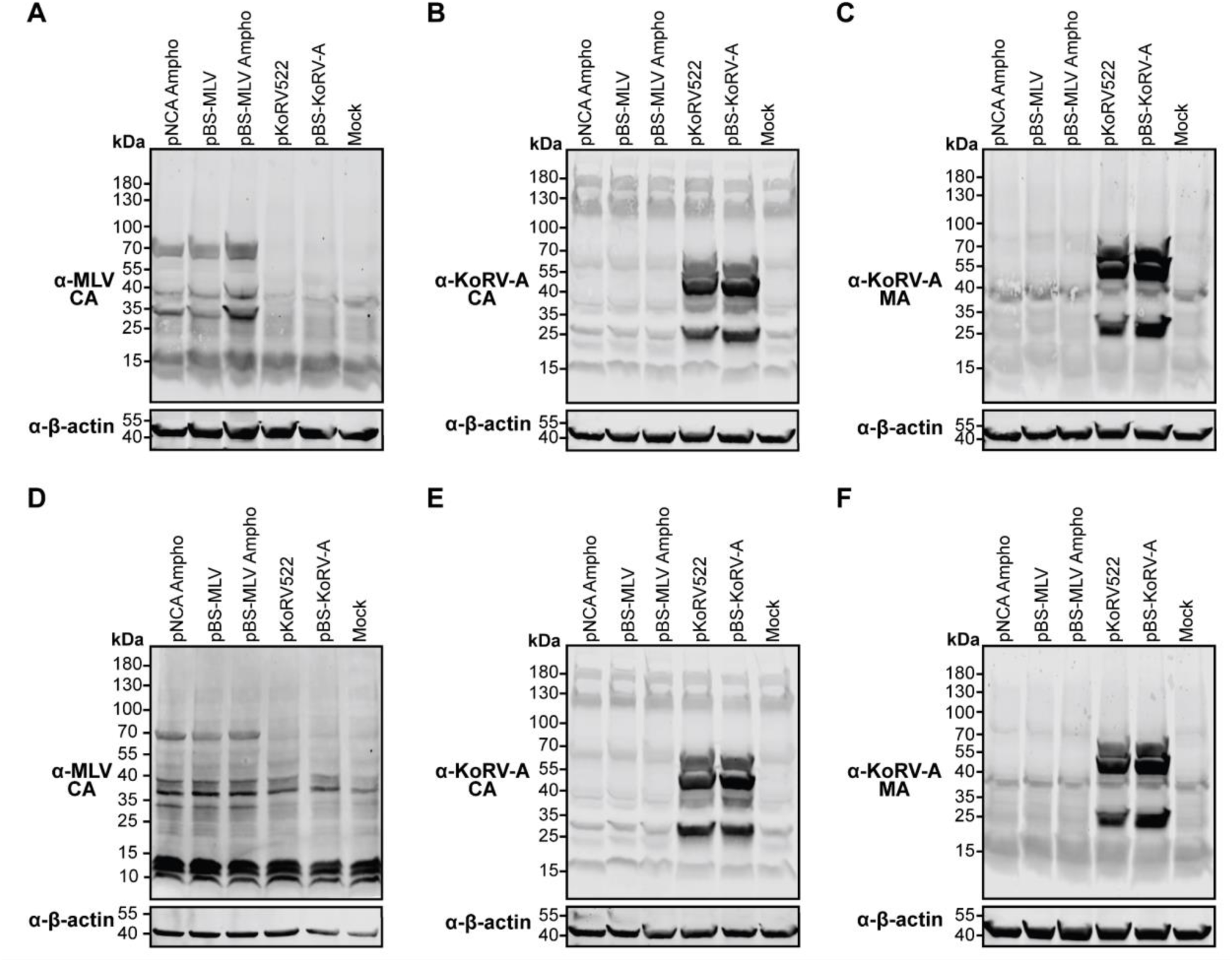
Additional biological replicates of Western blot analyses of transfected cells. HEK293T cells were transfected with the indicated proviral plasmids, and cell lysates were analyzed by Western blotting using antibodies against MLV CA (A, D), KoRV-A CA (B, E), or KoRV-A MA (C, F). β-Actin served as a loading control. Molecular masses are indicated in kDa.

**S5 Fig.**
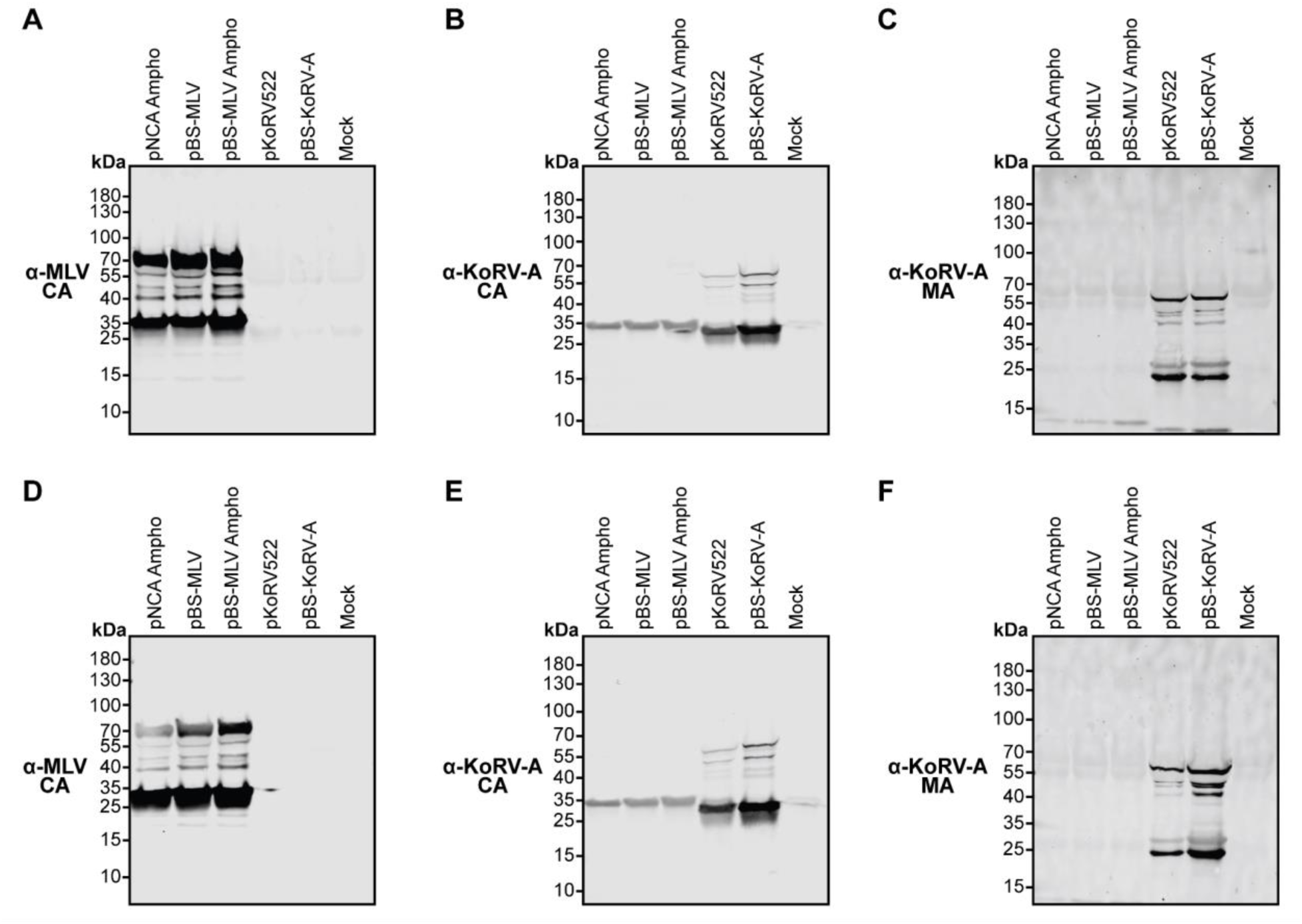
**Additional biological replicates of Western blot analyses of pelleted virions.**HEK293T cells were transfected with the indicated proviral plasmids. Virions were pelleted from equal volumes of culture supernatant and analyzed by Western blotting using antibodies against MLV CA (A, D), KoRV-A CA (B, E), or KoRV-A MA (C, F). Molecular masses are indicated in kDa.

**S6 Fig.**
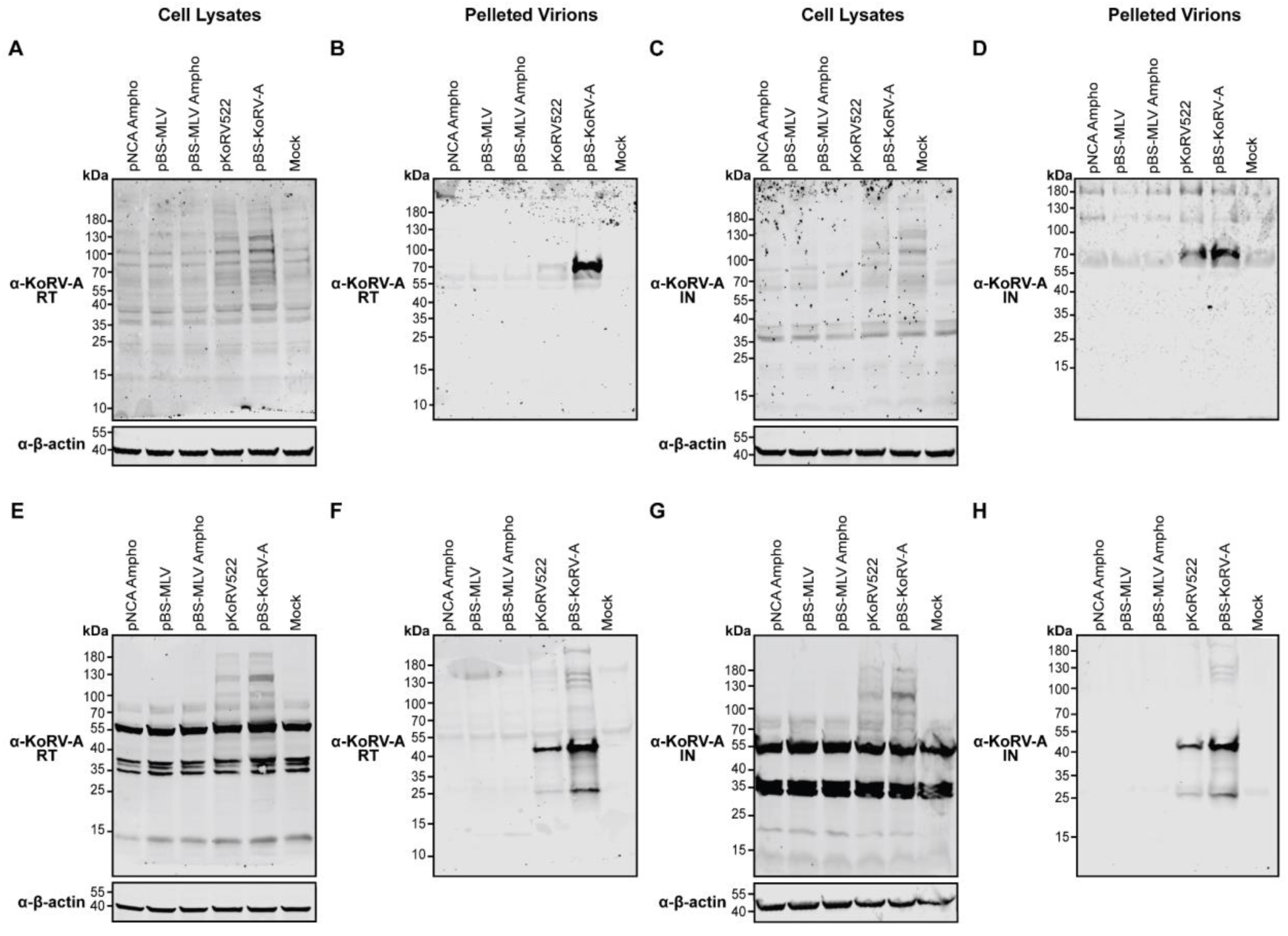
Additional biological replicates of Western blot analyses of KoRV-A RT and IN. Transfected HEK293T cell lysates (A, C, E, G) and virions pelleted from equal volumes of the corresponding culture supernatants (B, D, F, H) were analyzed by Western blotting. Samples were probed with antibodies specific for KoRV-A RT (A, B, E, F) or KoRV-A IN (C, D, G, H). β-Actin served as a loading control for the cell lysates. Molecular masses are indicated in kDa.

**S7 Fig.**
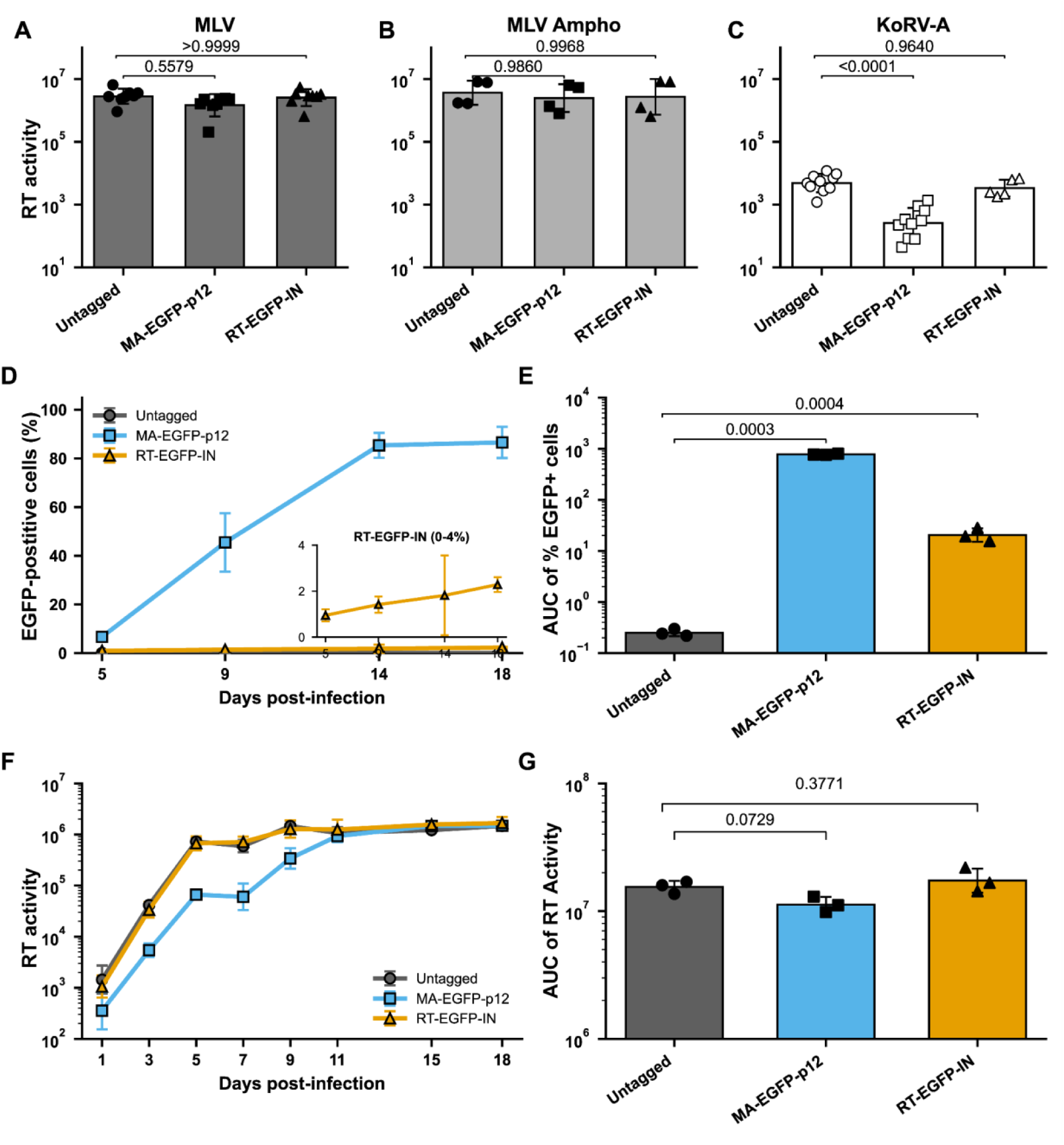
**Functional validation of EGFP-tagged retroviral reporter constructs.**(A–C) Exogenous RT activity in culture supernatants from HEK293T cells transfected with untagged, MA– EGFP–p12, or RT–EGFP–IN constructs in the MLV (A), MLV Ampho (B), or KoRV-A (C) background. Bars show geometric means ± geometric standard deviations, and points represent individual biological replicates. Statistical significance was evaluated using lognormal ordinary one-way ANOVA followed by Šídák’s multiple-comparisons test comparing each EGFP-tagged construct with the corresponding untagged construct. Exact Šídák-adjusted *p* values are shown. (D–G) HEK293T cells were challenged with cell-free virus supernatants generated from the indicated MLV Ampho constructs by spinoculation. The percentage of EGFP-positive cells (D) and exogenous RT activity in culture supernatants (F) were measured over time. The inset in (D) shows the RT–EGFP–IN data on an expanded 0–4% scale. Data in (D) are presented as means ± SD, whereas RT-activity data in (F) are presented as geometric means ± geometric standard deviations. (E, G) Area under the curve (AUC) for the percentage of EGFP-positive cells (E) and RT activity (G), calculated independently for each biological replicate using the trapezoidal rule. Bars show geometric means ± geometric standard deviations, and points represent individual biological replicates. AUC values were analyzed using repeated-measures ANOVA on log-transformed data, followed by planned paired comparisons between the untagged construct and each EGFP-tagged construct with Holm correction within each outcome. Exact Holm-adjusted *p* values are shown.

**S8 Fig.**
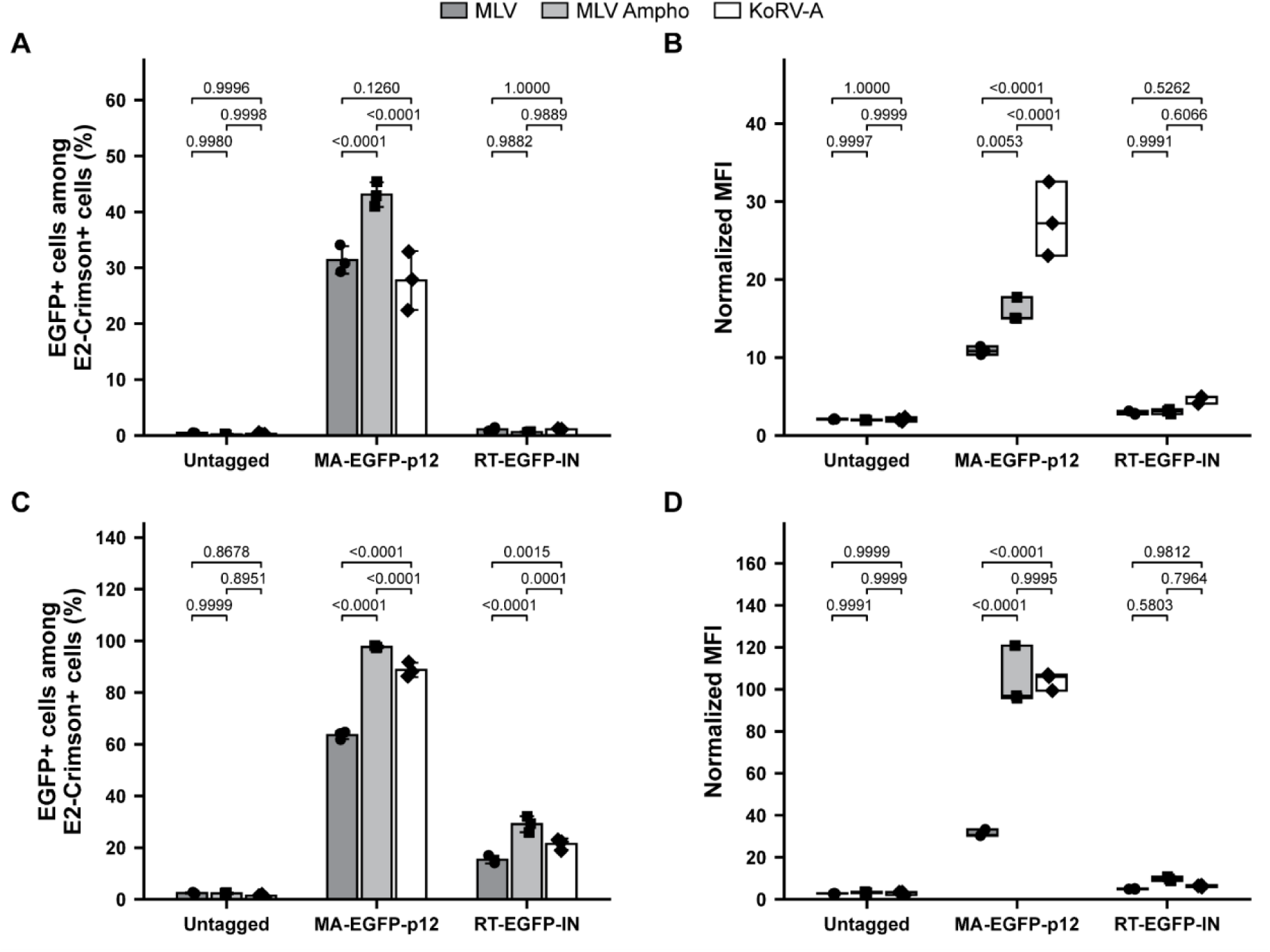
Additional independent experiments examining EGFP reporter expression in KoRV-A and MLV. (A, C) Percentage of E2-Crimson-positive cells that were also EGFP-positive following transfection with the indicated constructs in two independent experiments. Bars represent the mean ± SD of three parallel transfections, with individual values overlaid. (B, D) Normalized EGFP fluorescence, calculated as the ratio of EGFP MFI to E2-Crimson MFI within the EGFP-positive/E2-Crimson-positive population. Floating bars indicate the minimum and maximum values, horizontal lines indicate the medians, and individual values are overlaid. Statistical significance was evaluated separately for each experiment using ordinary two-way ANOVA with virus and reporter construct as factors, followed by Šídák-adjusted comparisons among viruses within each reporter construct. Exact adjusted *p* values are shown.

**S9 Fig.**
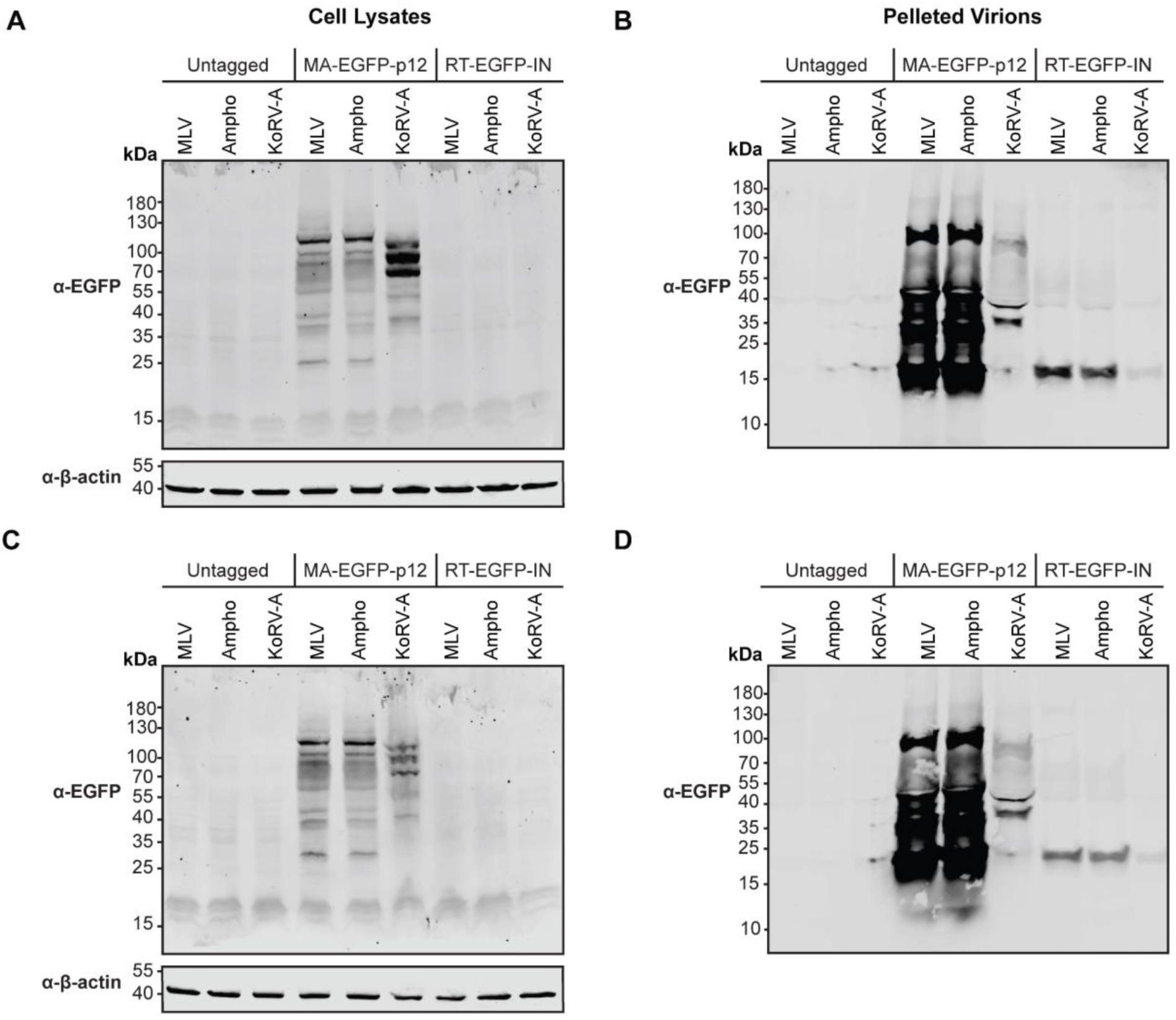
Additional biological replicates of Western blot analyses of EGFP-tagged viral constructs in transfected cells and pelleted virions. HEK293T cells were transfected with the indicated proviral plasmids. Transfected-cell lysates (A, C) and virions pelleted from equal volumes of culture supernatant (B, D) were analyzed by Western blotting using an antibody against EGFP. β-Actin served as a loading control for cell lysates. Molecular masses are indicated in kDa.

**S10 Fig.**
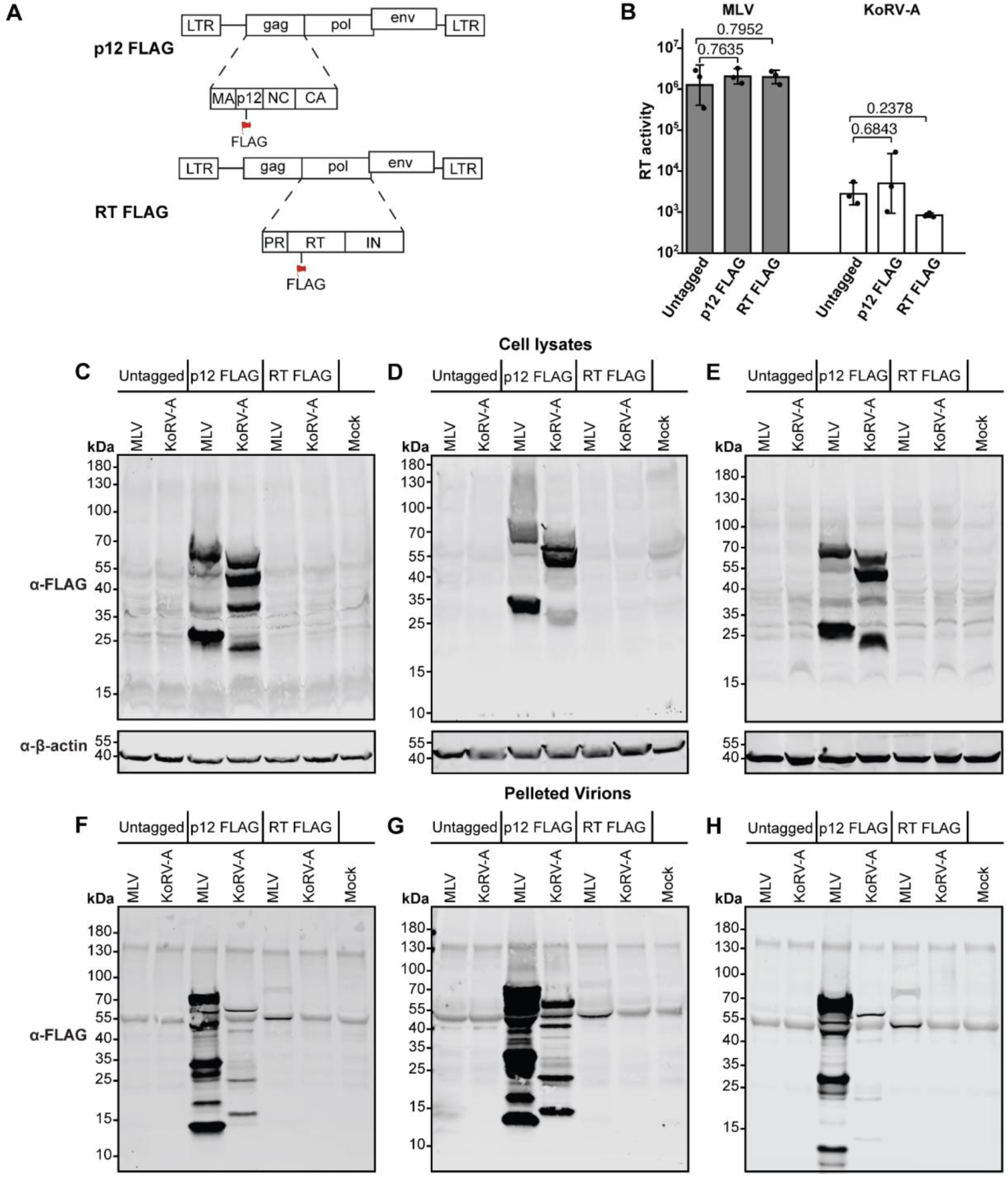
**Characterization of FLAG-tagged KoRV-A and MLV constructs.**(A) Schematic representation of the proviral constructs containing a FLAG epitope within p12 or RT in the KoRV-A and MLV backgrounds. (B) Exogenous RT activity in culture supernatants from HEK293T cells transfected with the corresponding untagged, p12-FLAG, or RT-FLAG proviral constructs. Data are presented as geometric means ± geometric standard deviations, with individual biological replicates shown as points. Statistical significance was evaluated by ordinary two-way ANOVA using log10-transformed values followed by Šídák’s multiple-comparisons test. Exact adjusted *p* values are shown. (C–E) Western blot analyses of transfected-cell lysates using an anti-FLAG antibody. β-Actin served as a loading control. (F–H) Western blot analyses of virions pelleted from equal volumes of the corresponding culture supernatants using an anti-FLAG antibody. Molecular masses are indicated in kDa.

**S11 Fig.**
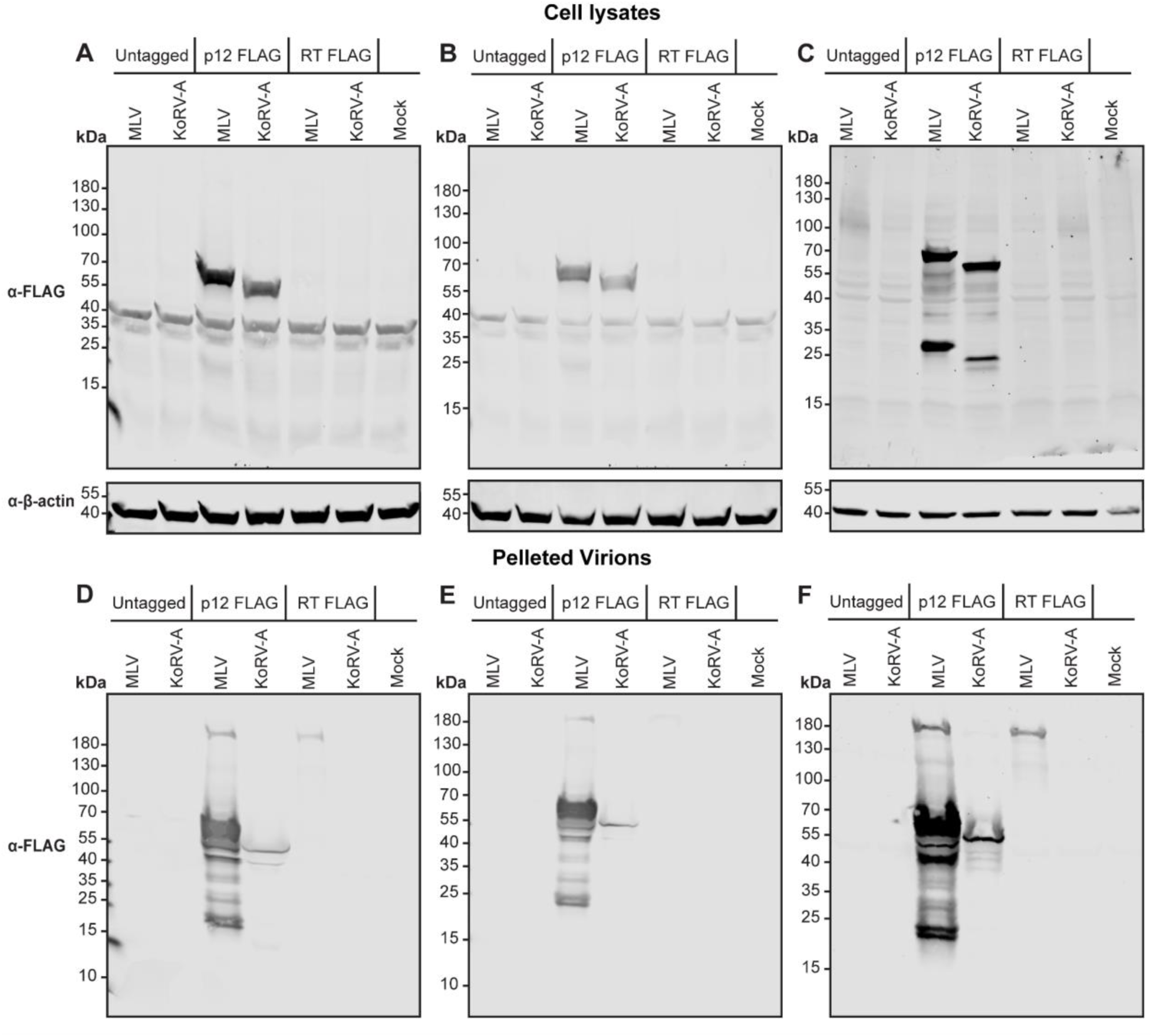
Expression and virion incorporation of FLAG-tagged PR mutants. (A–C) Western blot of transfected-cell lysates expressing the indicated FLAG-tagged PR-mutant constructs using an anti-FLAG antibody. β-Actin served as a loading control. (D–F) Western blot analyses of virions pelleted from equal volumes of the corresponding culture supernatants using an anti-FLAG antibody. Molecular masses are indicated in kDa.

**S12 Fig.**
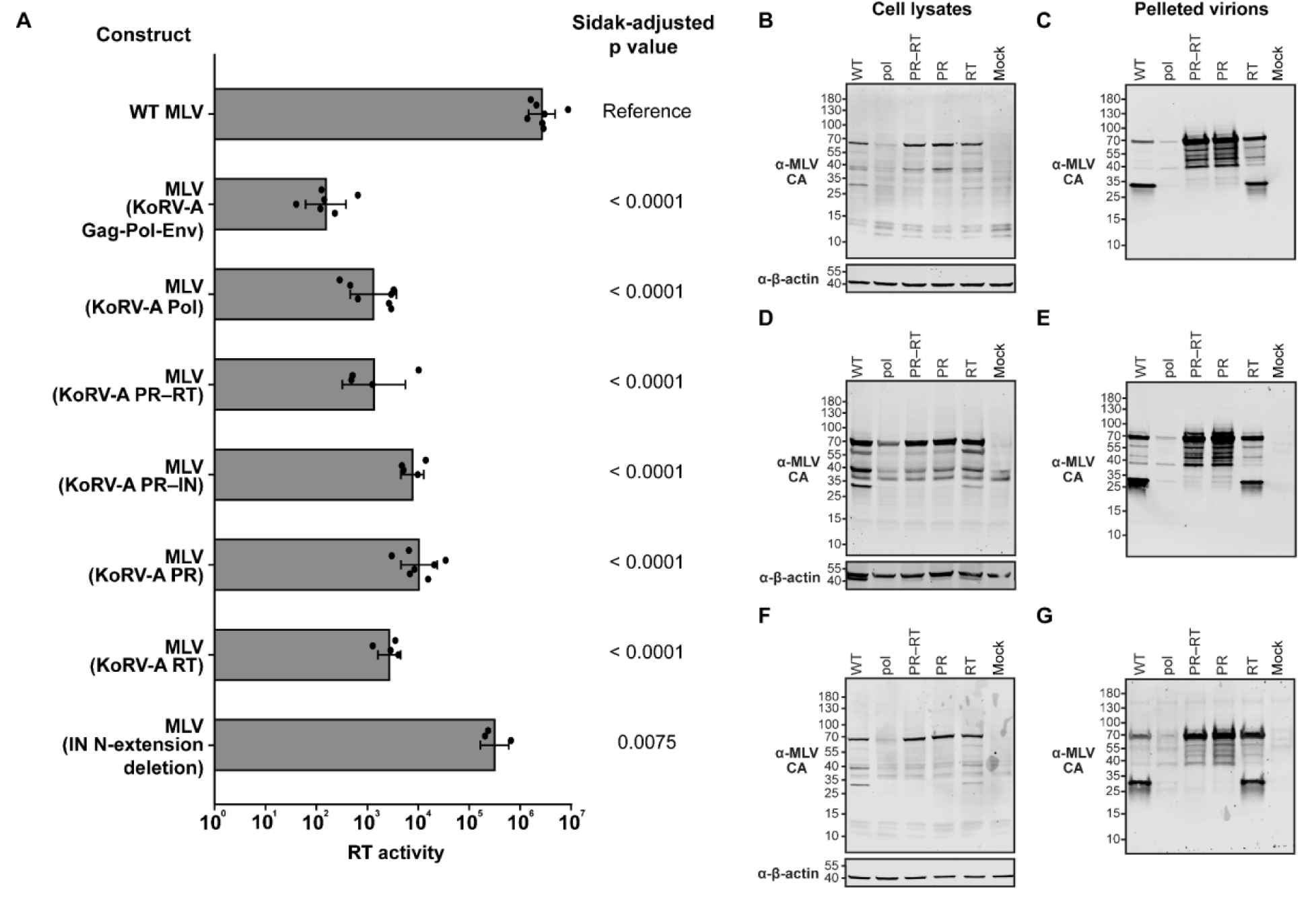
Characterization of chimeras generated in the MLV background. (A) Exogenous RT activity in culture supernatants from cells transfected with the indicated MLV chimeras. Data are presented as geometric means ± geometric standard deviations; individual points represent biological replicates. Statistical significance was evaluated using ordinary one-way ANOVA of log-transformed values followed by Šídák’s multiple-comparisons test comparing each chimera with WT MLV. Exact Šídák-adjusted *p* values are shown. (B–G) Western blot analysis of matched transfected-cell lysates (B, D, F) and virions pelleted from equal volumes of the corresponding culture supernatants (C, E, G) using an antibody against MLV CA. β-Actin served as a loading control for cell lysates. Molecular masses are indicated in kDa.

**S13 Fig.**
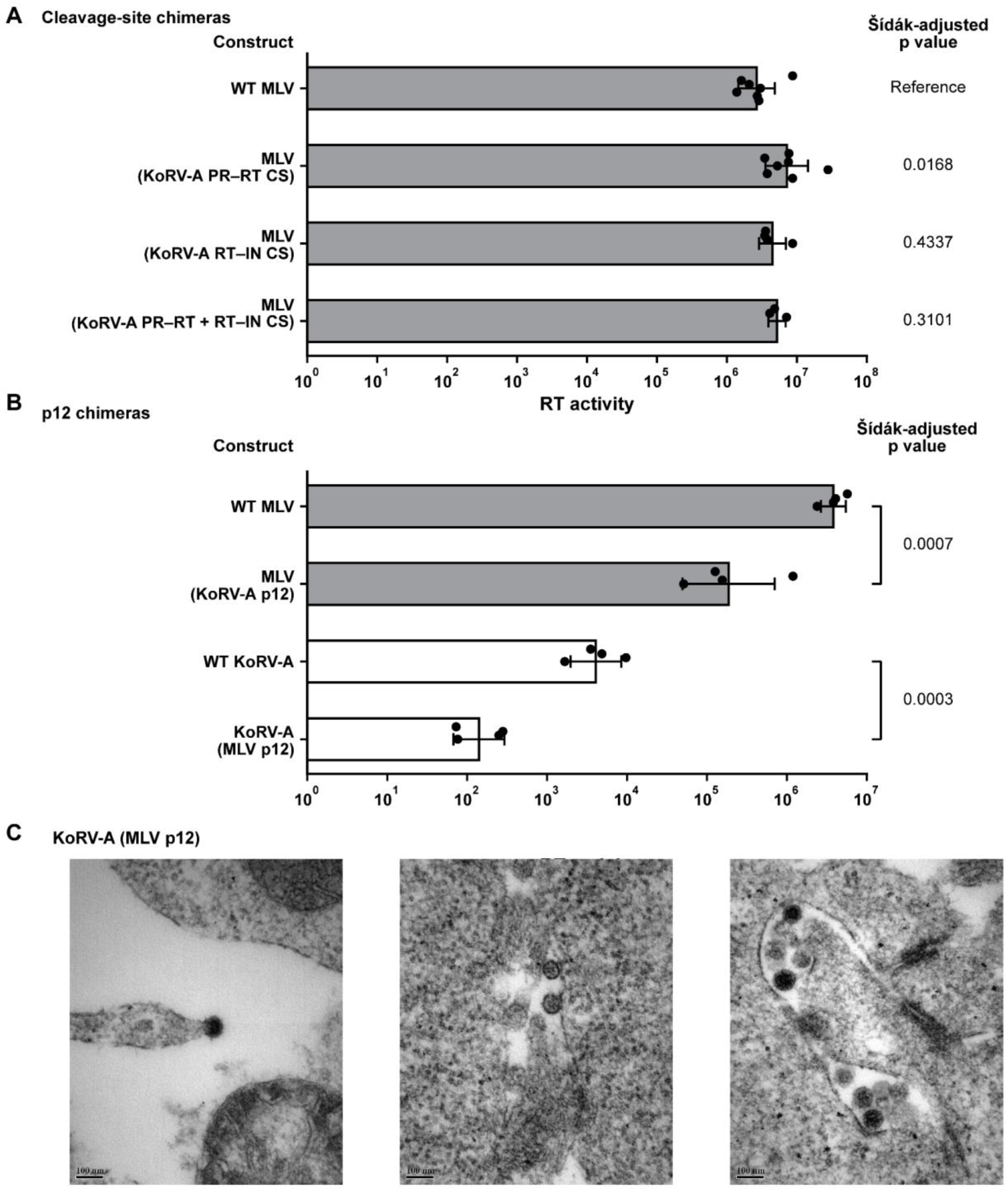
Characterization of MLV cleavage-site chimeras and reciprocal p12 chimeras. (A) Exogenous RT activity in culture supernatants from cells transfected with the indicated MLV chimeras containing KoRV-A cleavage-site sequences. (B) Exogenous RT activity of reciprocal p12 chimeras in which the p12 regions of KoRV-A and MLV were exchanged. (C) Representative TEM images of KoRV-A containing MLV p12. The scale bars represent 100 nm. RT activity data are presented as geometric means ± geometric standard deviations; individual points represent biological replicates. Statistical significance was evaluated using ordinary one-way ANOVA of log-transformed values followed by Šídák’s multiple-comparisons test comparing each chimera with its corresponding parental WT virus. Exact Šídák-adjusted *p* values are shown.

**S14 Fig.**
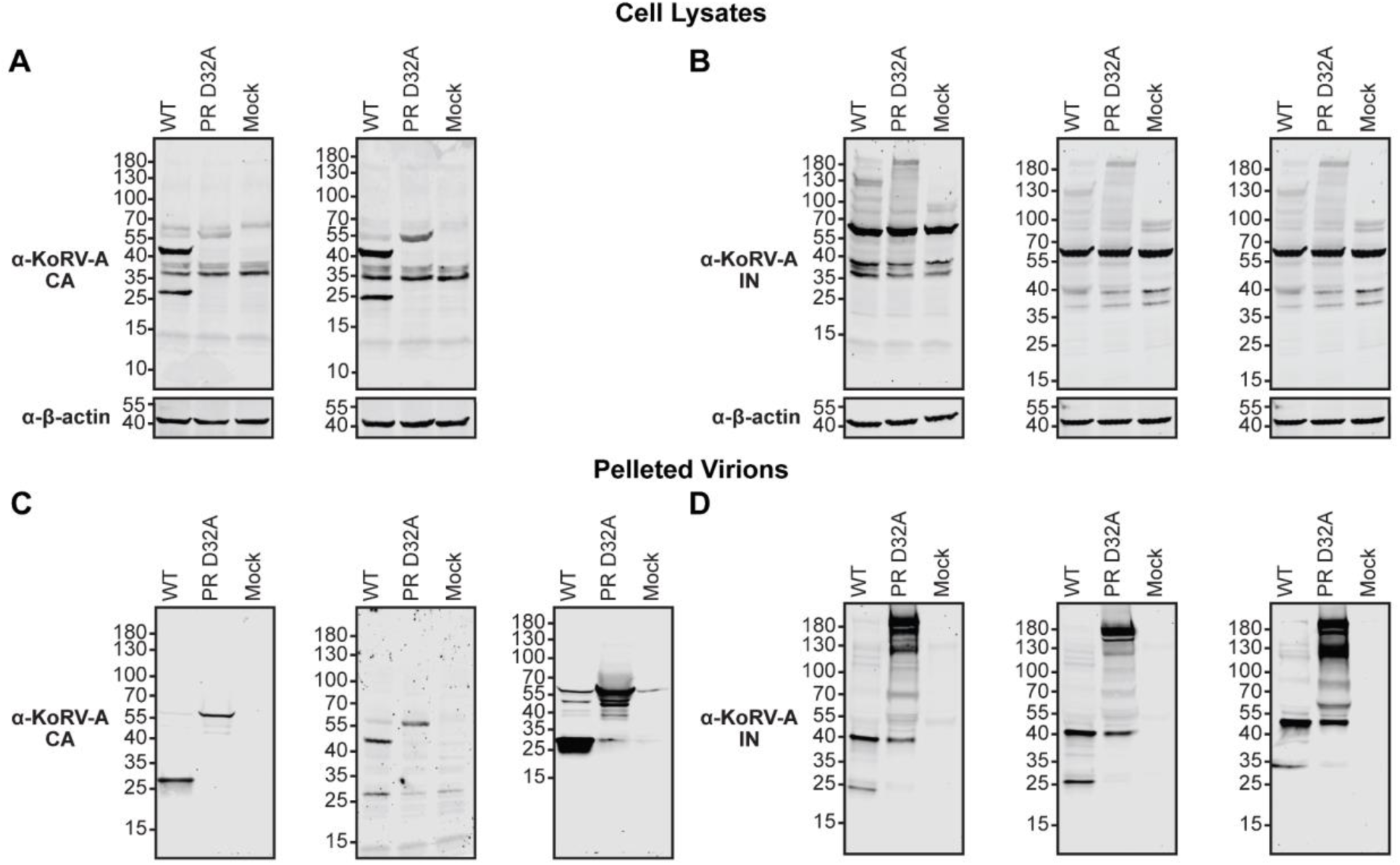
Western blot analysis of WT and PR-mutant KoRV-A in transfected cells and pelleted virions. (A, B) Western blot analysis of transfected-cell lysates from WT and PR-mutant KoRV-A using antibodies specific for KoRV-A CA (A; two independent biological replicates) or KoRV-A IN (B; three independent biological replicates). (C, D) Western blot analysis of virions pelleted from culture supernatants using antibodies specific for KoRV-A CA (C; three independent biological replicates) or KoRV-A IN (D; three independent biological replicates). Molecular masses are indicated in kDa.

**S15 Fig.**
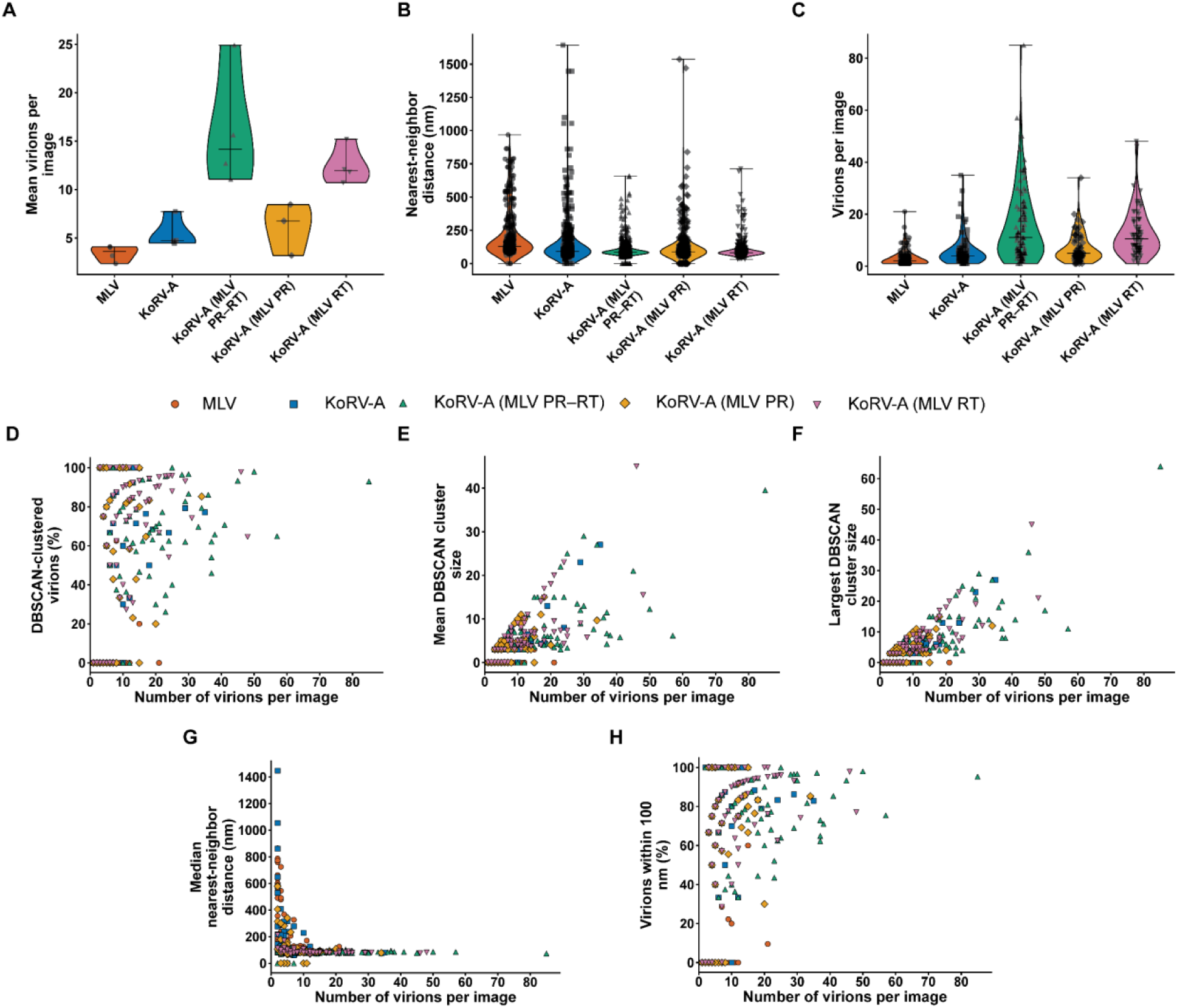
Virion abundance and its relationship with spatial clustering metrics for MLV, KoRV-A, and KoRV-A chimeras. (A) Mean number of virions per image for each biological replicate. Individual symbols represent biological replicates, and violin plots show the distribution of replicate-level values within each group. (B) Nearest-neighbor distances for individual virions. Each symbol represents one virion. This panel is descriptive and was not used for statistical testing because virions from the same image or biological replicate are not independent observations. (C) Number of virions detected in each analyzed image. Each symbol represents one image. (D–H) Relationships between the number of virions detected per image and (D) the percentage of virions assigned to a DBSCAN-defined cluster, (E) mean DBSCAN cluster size, (F) largest DBSCAN cluster size, (G) median nearest-neighbor distance, and (H) the percentage of virions with a nearest neighbor within 100 nm. Each symbol represents one image.

**S16 Fig.**
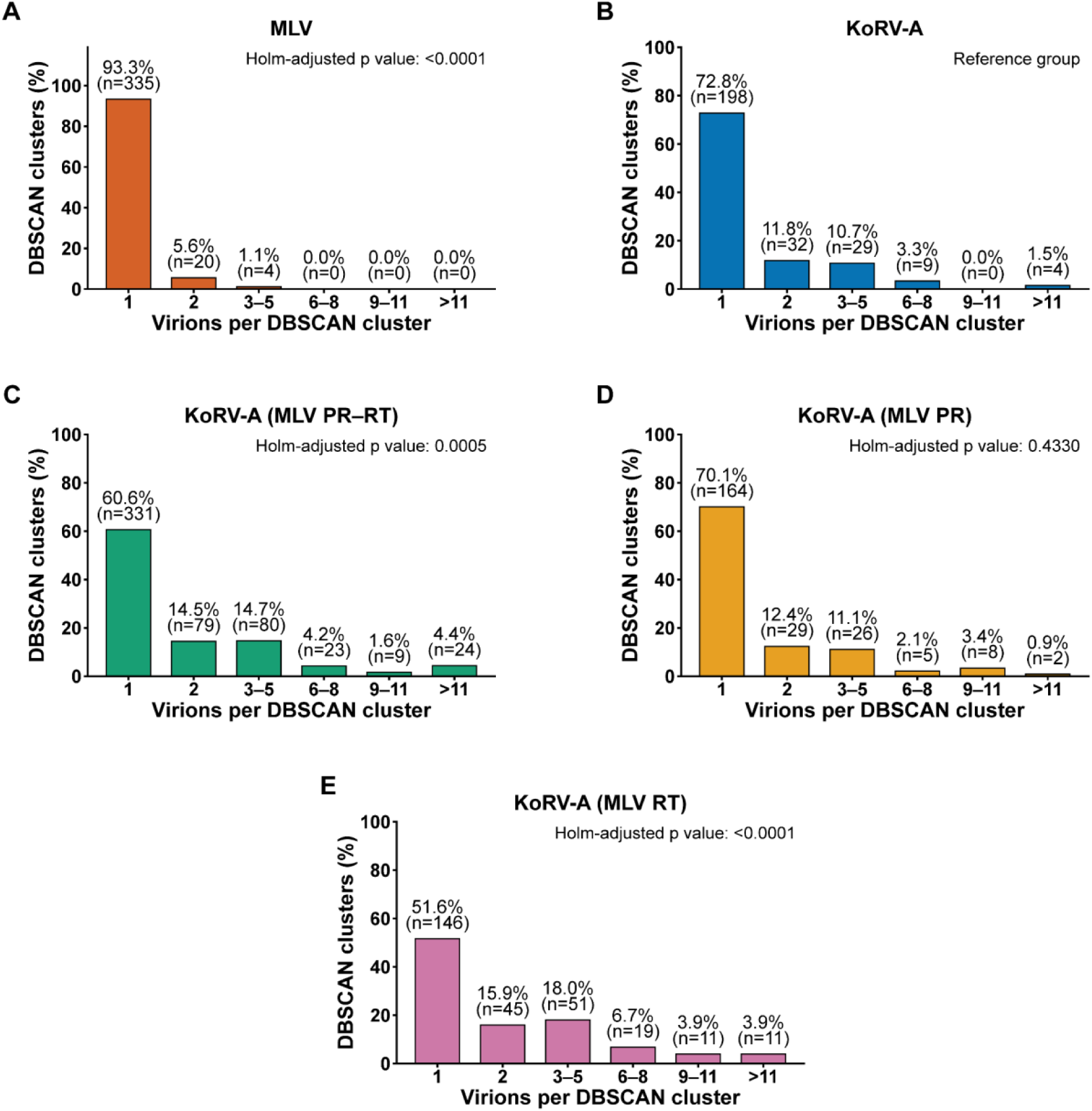
Distribution of extracellular particle-group sizes for MLV, KoRV-A, and KoRV-A chimeras. The distribution of particle-group sizes is shown for (A) MLV, (B) WT KoRV-A, (C) KoRV-A containing MLV PR–RT, (D) KoRV-A containing MLV PR, and (E) KoRV-A containing MLV RT. Particle groups were identified using DBSCAN with a 100-nm neighborhood radius and a minimum cluster size of one particle, permitting isolated virions to be classified as single-particle groups. Groups were categorized as containing 1, 2, 3–5, 6–8, 9–11, or >11 virions. Bars indicate the percentage of particle groups assigned to each size category; percentages and corresponding numbers of particle groups (*n*) are shown above the bars. Pooled, unbinned particle-group sizes were compared between WT KoRV-A and each of the other groups using two-sided Mann–Whitney U tests, with Holm correction across the four comparisons. Exact Holm-adjusted *p* values are shown.

**S17 Fig.**
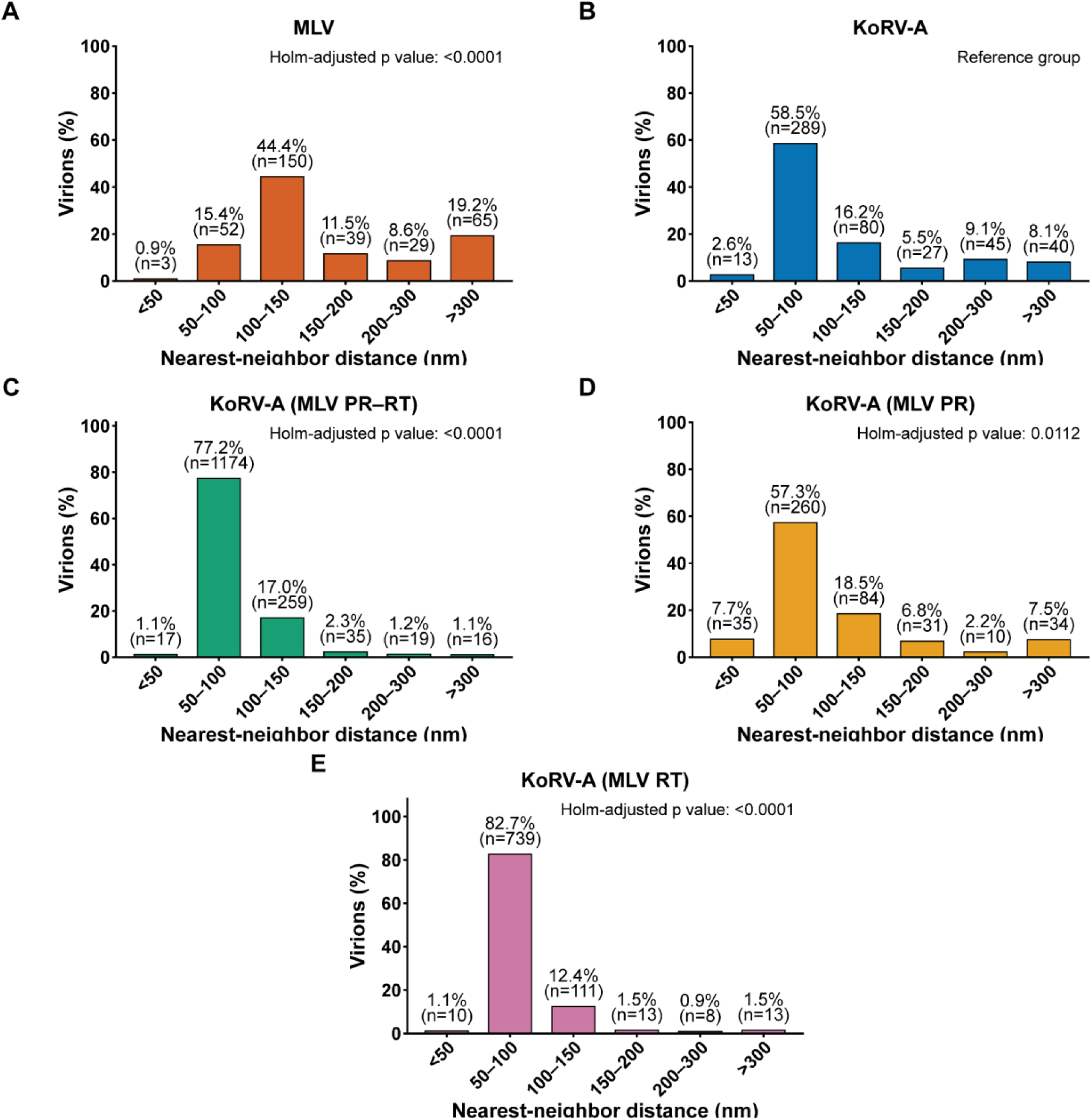
Distribution of virion nearest-neighbor distances. Nearest-neighbor-distance distributions are shown for (A) MLV, (B) KoRV-A, (C) KoRV-A containing MLV PR–RT, (D) KoRV-A containing MLV PR, and (E) KoRV-A containing MLV RT. For each virion, the distance to its nearest neighboring virion within the same TEM image was calculated and assigned to the indicated distance interval. Bars show the percentage of virions in each interval; percentages and numbers of virions (*n*) are indicated above the bars. Virions in images containing only one particle were excluded because a nearest-neighbor distance could not be calculated. Pooled, unbinned nearest-neighbor distances were compared between KoRV-A and each of the other groups using two-sided Mann–Whitney U tests, with Holm correction across the four comparisons. Exact Holm-adjusted *p* values are shown.

**S18 Fig.**
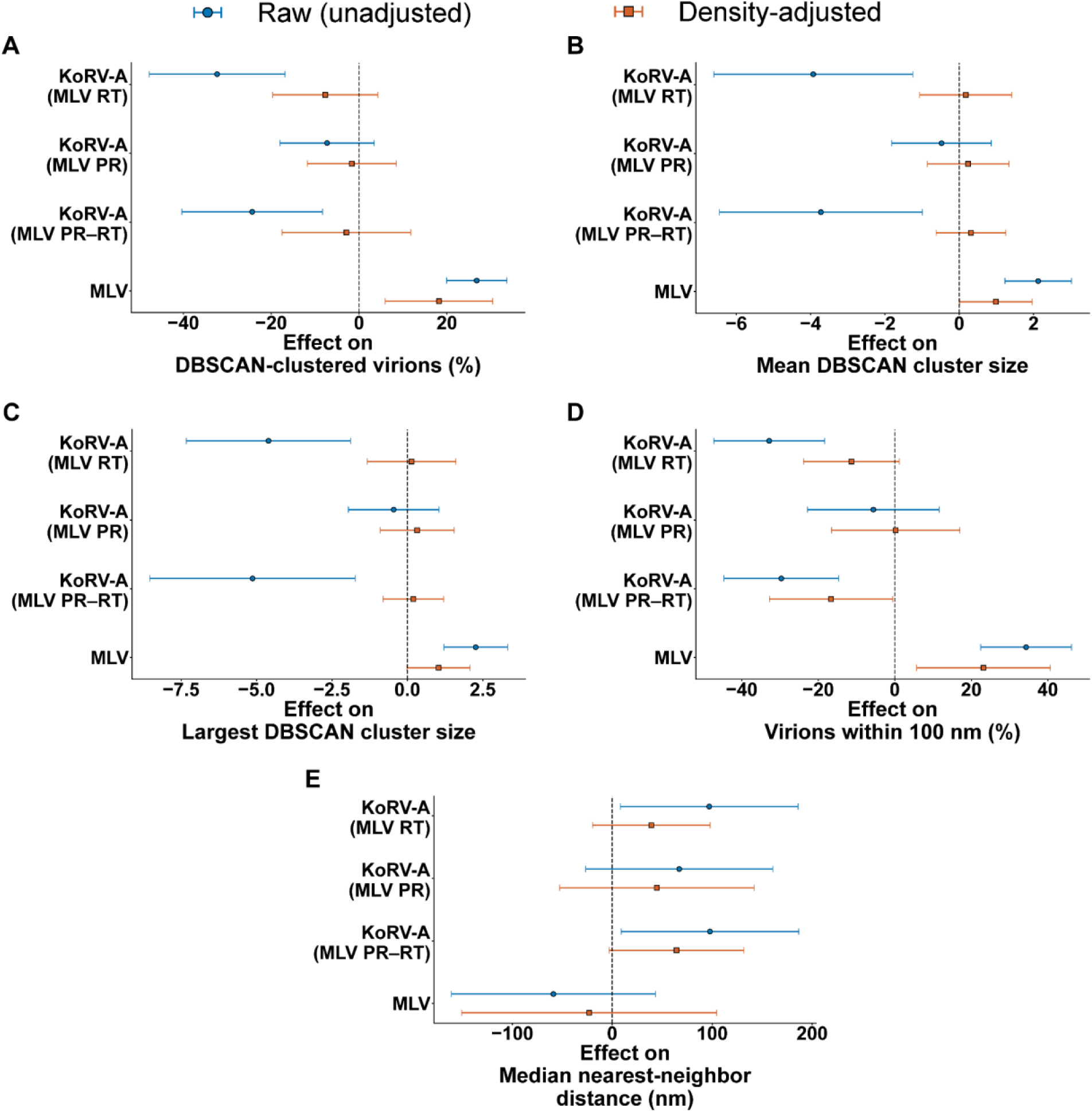
Particle-density-adjusted analysis of extracellular virion organization. (A–E) Raw and particle-density-adjusted differences between KoRV-A and each indicated comparison group for (A) the percentage of virions assigned to a DBSCAN-defined cluster, (B) mean DBSCAN cluster size, (C) largest DBSCAN cluster size, (D) the percentage of virions with a nearest neighbor within 100 nm, and (E) median nearest-neighbor distance. Blue circles indicate unadjusted group effects, and orange squares indicate effects after adjustment for the natural logarithm of the number of virions detected per image. Points represent regression coefficients calculated as KoRV-A minus the indicated comparison group; error bars indicate 95% confidence intervals. The vertical dashed line denotes no difference between groups. Models were fitted at the image level, with standard errors clustered by biological replicate.

